# Exploration of Orally Disintegrating Tablet for Sublingual Vaccination against Mucosal Bacterial Infection

**DOI:** 10.64898/2026.03.14.711758

**Authors:** Yitong Liu, Qinming Cai, Xiaoming Hu, Xin Liu, Peilin Guo, Yu Zhang, Hongyi Liu, Wenjing Wang, Diwei Zheng, Chao Pan, Lijia Guo, Xijiao Yu, Qihong Zhang, Dongfang Wang, Yefeng Qiu, Dongshu Wang, Wenxian Li, Yi Du, Guanghui Ma, Junji Xu, Shuang Wang, Yi Liu, Wei Wei

## Abstract

Oral mucosal bacterial infections impose a substantial global disease burden, yet current clinical management typically reduces microbial load only transiently and rarely establishes durable protection at the oral surface. Analysis of 200 patients with periodontitis revealed that elevated levels of pathogen-specific salivary secretory Immunoglobulin A (sIgA) were strongly associated with reduced bacterial burden and improved clinical periodontal outcomes, identifying sIgA as a key determinant of effective oral protection. Guided by this observation, we developed a sublingual, orally disintegrating tablet vaccine (Capot) that incorporates bacterial extracellular vesicles providing a comprehensive repertoire of native antigens and multiple pathogen-associated molecular patterns, encapsulated within a calcium phosphate nanoshell to enable safe transmucosal delivery to submandibular lymph nodes. The rapidly disintegrating tablet format minimizes inadvertent swallowing and enhances local mucosal bioavailability. In mice and non-human primates, Capot induced robust and long-lasting salivary sIgA responses without overt oral mucosal or gastrointestinal inflammation and conferred strong protection against primary, recurrent, and antibiotic-resistant periodontitis. Together, these findings establish sublingual tablet vaccination as a practical strategy for selectively engaging oral mucosal immunity and preventing chronic bacterial diseases at oral mucosal surfaces.

## Introduction

Oral infectious diseases constitute a widespread and underappreciated global health burden, affecting billions of individuals worldwide^1,2,3^. Persistent bacterial colonization of the oral cavity can lead to chronic inflammation, tissue destruction, pain, and diminished quality of life^1,4^. As one of the representative diseases, periodontitis is a highly prevalent inflammatory disorder of the tooth-supporting structures driven by pathogenic oral bacteria such as *Porphyromonas gingivalis* (*P.g*) ^5,6,7^. It remains a leading cause of tooth loss and has been increasingly associated with systemic disorders, including cardiovascular disease^8^, diabetes^9^, and adverse pregnancy outcomes^10^. Current clinical management relies primarily on mechanical debridement and antibiotic therapy, which can transiently reduce microbial burden but rarely confer durable protection at the oral surface ^11,12,13^. Moreover, repeated antibiotic exposure contributes to microbial imbalance and accelerates the emergence of drug-resistant strains, underscoring the urgent need for alternative preventive strategies^2,14,15^.

With the advancement of immunology, researchers have realized that saliva constitutes the first line of immune defense in the oral cavity and functions as an immunologically active fluid^2,16,17^. Beyond mechanical clearance, saliva contains immunoglobulins, predominantly Immunoglobulin G (IgG) and secretory Immunoglobulin A (sIgA), that restrict bacterial adherence and invasion through immune exclusion and neutralization at mucosal surfaces^18^. Despite this recognized importance, the relative contribution of individual salivary antibody classes to protection against oral bacterial disease, as well as the relevance of circulating serum antibodies, has remained unclear. Using periodontitis as a clinically relevant starting point, we analyzed saliva and serum from 200 individuals with distinct periodontal outcomes and found that high levels of pathogen-specific salivary sIgA, but not salivary IgG or serum IgG/IgA, were strongly associated with favorable prognosis. These findings identify salivary sIgA as a central determinant of effective oral protection and provide a rationale for immunization strategies that selectively induce localized mucosal sIgA responses in the oral cavity.

We envision that, in contrast to parenteral vaccination that predominantly elicits systemic immunity, sublingual immunization is uniquely positioned to drive targeted mucosal antibody responses^19^. The sublingual mucosa represents an anatomically favorable site for engaging oral mucosal immune circuits. Antigens delivered beneath the tongue may access a dense lymphatic network that drains directly to the submandibular lymph nodes (sLNs), a central inductive hub for oral immune responses^20,21^. Engagement of this pathway promotes the generation of IgA-secreting plasma cells, enabling localized antibody production within the oral cavity^20^. However, despite its theoretical promise, the development of safe and effective sublingual vaccines remains constrained by several interrelated challenges. Vaccine particles must be appropriately sized to enable lymphatic drainage to the sLNs, as micron-sized formulations fail to reach this site^22,23,24^. In addition, potent mucosal adjuvants (often pattern recognition receptor agonists) can trigger strong innate signaling in oral epithelial/structural cells, leading to chemokine/cytokine release and recruitment of inflammatory cells, which may manifest as local reactogenicity such as irritation and inflammation^25^. Third, the highly dynamic oral environment promotes rapid clearance and swallowing of administered formulations, limiting mucosal exposure and increasing the risk of off-target gastrointestinal effects. Together, these barriers have hindered the translation of sublingual vaccination into clinically practical modalities.

To overcome these challenges, we developed a sublingual tablet-based oral mucosal vaccine platform designed to coordinate immunogenicity, safety, and delivery efficiency. This system integrates bacteria-derived extracellular nanovesicles coated with a pH-responsive calcium phosphate (CaP) nanoshell and formulates them into a rapidly disintegrating sublingual tablet, termed Capot. Upon administration, the tablet rapidly disintegrates in saliva, prolonging residence time beneath the tongue and minimizing unintended swallowing. The CaP coating shields immunostimulatory motifs during mucosal transit, enabling the nanovesicles to safely penetrate the sublingual mucosa, reach the sLNs, and be efficiently taken up by antigen-presenting cells. This targeted delivery results in robust activation of the oral mucosal immune compartment, inducing potent salivary sIgA responses and enhancing antibacterial activity. The platform demonstrates consistent efficacy across multiple disease contexts, including primary, recurrent, and antibiotic-resistant periodontitis, in both murine and non-human primate models, supporting its potential as a broadly applicable strategy for oral mucosal vaccination.

## Results

### Association of elevated salivary *P.g*-specific sIgA with periodontal outcomes

Given the unclear role of individual serum and salivary antibody classes in protection against oral bacterial diseases, we used periodontitis as a clinically relevant model and enrolled 200 patients with periodontitis who received primary periodontal therapy aimed at eliminating *P.g* from the oral cavity (Figure 1A). Patients were re-evaluated three months after treatment. During this period, all participants were naturally re-exposed to *P.g* bacteria through their normal daily diets. At the follow-up visit, we assessed changes in periodontal pocket probing depth (PD) and bleeding on probing (BOP) as clinical indicators of periodontal outcomes. Meanwhile, both serum and saliva were collected to quantify *P.g*-specific IgG and IgA titers in serum, as well as *P.g*-specific IgG and sIgA titers in saliva. In addition, we analyzed salivary *P.g* bacterial abundance using 16S rDNA sequencing and quantitative PCR (qPCR).

**Figure 1.**
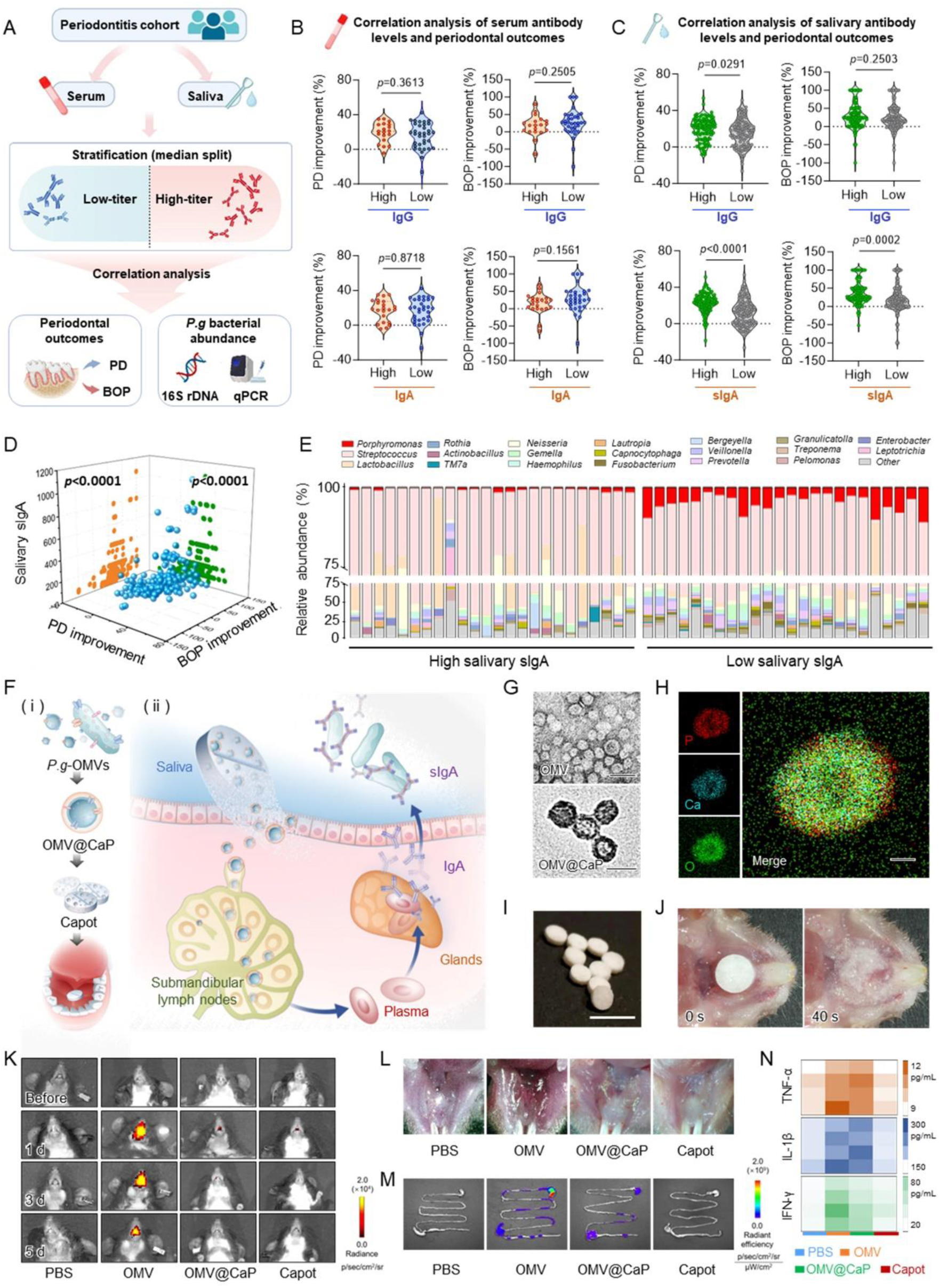
Clinical evidence supporting the dominant role of salivary *P.g*-specific sIgA antibody in periodontal outcomes and the design of the Capot sublingual oral mucosal vaccine platform. A) Schematic illustrating the clinical analysis used to evaluate the association between antibody classes and periodontal outcomes. PD = periodontal pocket probing depth; BOP = bleeding on probing. B) Subgroup analysis comparing periodontal outcomes between patients with high versus low serum *P.g*-specific antibody levels (median split; high IgG group, n = 19, low IgG group, n = 35; high IgA group, n = 22, low IgA group, n = 32). C) Subgroup analysis comparing periodontal outcomes between patients with high versus low salivary *P.g*-specific antibody levels (median split; high IgG group, n = 102, low IgG group, n = 98; high sIgA group, n = 96, low sIgA group, n = 104). D) Pearson correlation analyses examining relationships between PD improvement, BOP improvement, and salivary *P.g*-specific sIgA antibody titers (n = 200). Blue dots indicate individual patients; green and orange projections represent PD-sIgA and BOP-sIgA correlations, respectively. E) 16S rRNA gene sequencing analysis of salivary microbiota in family level from representative patients with high or low salivary *P.g*-specific sIgA levels (n=24). F) Schematic of the Capot design (i) and its action mechanism (ii). G) Transmission electron microscopy (TEM) images of *P.g*-OMV (top) and OMV@CaP (bottom). Scale bar = 100 nm. H) Elemental mapping of OMV@CaP confirming CaP coating. Scale bar = 20 nm. I) Representative photograph of Capot. Scale bar = 4 mm. J) Photographs showing Capot disintegration in the mouse oral cavity. K) Representative bioluminescence images of IFN-γ-IRES-Venus-Akaluc reporter mice before and after sublingual administration of PBS, OMV, OMV@CaP, or Capot at 1 d, 3 d, and 5 d. L) Representative photographs of sublingual mucosa at 24 h after administration of the indicated formulations. M) Representative fluorescence images of gastrointestinal tissues at 24 h after sublingual administration of PBS, Cy7-labelled OMV, OMV@CaP, or Capot. N) Concentrations of TNF-α, IL-1β, and IFN-γ in gastrointestinal tissues at 24 h after administration of PBS, OMV, OMV@CaP, or Capot. Data in (B, C) are shown as violin plots. Statistical significance was tested with unpaired two-tailed t-tests (B, C) and Pearson correlation analysis (D). *p* values have been presented in the figure.

To evaluate the relationship between antibody titers and periodontal outcomes, we first stratified patients into high- and low-titer groups for each antibody type (serum IgG, serum IgA, salivary IgG, and salivary sIgA), using the median titer across all patients as the threshold. We then compared PD and BOP improvements within each corresponding subgroup (Figure 1B and 1C). Among these, salivary sIgA subgroup demonstrated the highest association. Patients with higher *P.g*-specific salivary sIgA titers showed greater improvements in both PD and BOP compared to those with lower titers. Correlation analyses further supported this association, revealing a positive relationship between salivary sIgA titers and both clinical indicators (Figure 1D and Figure S1A). To integrate all four antibody classes and their relationship to periodontal outcomes, we constructed a predictive model using XGBoost. SHAP (SHapley Additive exPlanations) analysis of the model ranked salivary *P.g*-specific sIgA as the most influential feature for both PD and BOP improvements (Figure S1B). Based on these findings, we further analyzed salivary *P.g* bacterial abundance using 16S rDNA sequencing and qPCR, which revealed that patients with higher salivary sIgA titers had significantly lower salivary *P.g* abundance than those with lower titers (Figure 1E and Figure S1C). Above data collectedly supporting a dominating antibacterial role for *P.g*-specific sIgA in the oral environment.

### Design and construction of the Capot sublingual vaccine

Given the observed association between salivary *P.g*-specific sIgA and clinical periodontal outcomes, and the capacity of sublingual immunization to deliver antigens to sLNs and induce mucosal IgA responses, we sought to develop a sublingual vaccine to enhance salivary sIgA levels and reduce periodontal *P.g* burden. As the vaccine core, we selected *P.g*-derived outer membrane vesicles (OMVs), which intrinsically present a broad antigen repertoire along with multiple pathogen-associated molecular patterns (PAMPs), enabling multi-antigen immune activation^29^. Their nanoscale size further facilitates passive lymphatic trafficking to sLNs following sublingual delivery^30^. However, direct sublingual administration of OMVs can induce local mucosal irritation and inflammation, as well as gastrointestinal inflammation resulting from vesicle swallowing. To mitigate these limitations while preserving immunogenicity, we coated OMVs with a biocompatible, pH-responsive CaP shell *via* biomimetic mineralization and formulated them into fast-disintegrating sublingual tablet, termed Capot (Figure 1F), a design intended to mask surface PAMPs during mucosal transit, reduce local and gastrointestinal irritation, and improve sublingual bioavailability.

Transmission electron microscopy, elemental mapping and dynamic light scattering together confirmed that CaP coating produced uniform core-shell OMV@CaP nanoparticles, increasing the average diameter from 50.75 nm to 105.7 nm (Figure 1G,H; Figure S2A,B). Thermogravimetric and differential scanning calorimetry analyses estimated a CaP mass fraction of approximately 50% (Figure S2C). SDS-PAGE analysis further showed that protein banding patterns of OMV cores were preserved relative to naked OMVs (Figure S2D), indicating successful encapsulation. For sublingual delivery, OMV@CaP particles were lyophilized and formulated into fast-disintegrating tablets (Capot) (Figure 1I). Upon placement beneath the murine tongue, Capot rapidly disintegrated and fully dissolved within 40 seconds (Figure 1J), a property expected to enhance mucosal contact while minimizing swallowing. Capot also exhibited strong storage stability. After six months storage at room temperature, the tablets retained structural integrity and appearance without detectable deconstruction (Figure S2E). Notably, OMV@CaP particles released from stored tablets retained much similar size distribution, surface charge, and dendritic cell (DC)-activating capacity (Figure S2F-H), laying the foundation for efficient *in vivo* sLN accumulation and subsequent mucosal immune activation after long-term storage at room temperature.

### Attenuation of oral and gastrointestinal inflammation by the Capot formulation

Having established the OMV@CaP nanoparticle formulation and its integration into fast-disintegrating Capot tablet, we next evaluated whether this design effectively mitigates oral and gastrointestinal inflammatory responses, consistent with the underlying formulation rationale. To assess local inflammatory signaling, IFN-γ-IRES-Venus-AkaLuciferase reporter mice were sublingually administered free OMVs, OMV@CaP in solution, or Capot tablet and monitored by longitudinal bioluminescence imaging over five days. Free OMVs induced strong and sustained IFN-γ signals at the oral cavity, whereas both OMV@CaP and Capot treatments resulted in markedly attenuated signal intensity throughout the observation period (Figure 1K and S3A). Consistently, local cytokine profiling revealed reduced levels of TNF-α, IL-1β, and IL-6 in OMV@CaP- and Capot-treated mice compared to the free OMV group (Figure S3B). Macroscopically, sublingual mucosal erythema was evident in the free OMV group but was absent in both OMV@CaP-and Capot-treated mice (Figure 1L). Histological examination further confirmed substantially reduced inflammatory cell infiltration in tongue tissues from these groups (Figure S3C). Collectively, these findings demonstrate that CaP encapsulation effectively dampens local inflammatory responses, likely by masking surface-exposed PAMPs on OMVs during transmucosal transport^31^.

We next examined whether formulation into a fast-disintegrating tablet conferred additional protection against swallowing-associated gastrointestinal exposure beyond CaP encapsulation alone. Cy7-labeled vesicle formulations were tracked after sublingual administration. *In vivo* imaging showed no detectable Cy7 signal in the gastrointestinal tract of Capot-treated mice, in contrast to clear signals observed in the OMV@CaP solution group (Figure 1M). Correspondingly, intestinal levels of inflammatory cytokines in the Capot group were significantly reduced and approached those observed in PBS-treated controls (Figure 1N and Figure S3D). This reduction in gastrointestinal adverse effects is attributable to the fast-disintegrating tablet format, which reduces free saliva volume and increases local viscosity, thereby prolonging sublingual residence time and minimizing inadvertent swallowing. In addition, no abnormalities in hematological parameters and blood biochemistry were detected following Capot administration (Figure S3E). Together, these data support the favorable safety profile of Capot for sublingual vaccination and validate its design for localized mucosal immune engagement with minimal off-target inflammation.

### Lymphatic accumulation and immune cell activation induced by Capot

After confirming the *in vivo* safety of Capot, we next examined its accumulation in sLNs following sublingual vaccination. OMVs were labeled with Cy7 and further formulated into Cy7-labeled OMV@CaP and Capot. As shown in Figure 2A and Figure S4A, both OMV@CaP and Capot accumulated in sLNs, with Capot exhibiting superior enrichment, consistent with reduced swallowing and improved oral mucosal bioavailability conferred by the disintegrating tablet format. Within sLNs, OMV@CaP nanoparticles were internalized by DCs: the proportion of Cy7⁺ DCs reached 72.8% in the Capot group, markedly higher than the 48.3% observed with OMV@CaP (Figure 2B). Following uptake, the CaP shell underwent acid-triggered dissolution within DC lysosomes (Figure S4B), enabling controlled intracellular release of vesicle antigens. Consistent with above data, DC activation and antigen-presentation markers (CD40, CD80, and MHC II) were more robustly upregulated in the Capot group (Figure 2C and Figure S4C). This enhancement of DC further promoted robust CD4 T-cell activation in sLNs, with Capot inducing the highest expression of CD25 and Ki67 among all groups (Figure 2D). To further assess CD4⁺ T cell-B cell interactions, cleared sLN tissues were labeled with CD4 and B220 antibodies and analyzed by light-sheet microscopy. As shown in Figure 2E and Figure S4D-E, Capot induced pronounced B-cell follicle expansion compared with PBS and OMV@CaP. Moreover, CD4 T-cell infiltration into B-cell zones was increased 6.7-fold relative to PBS control, indicating enhanced CD4 T-B cell collaboration.

**Figure 2.**
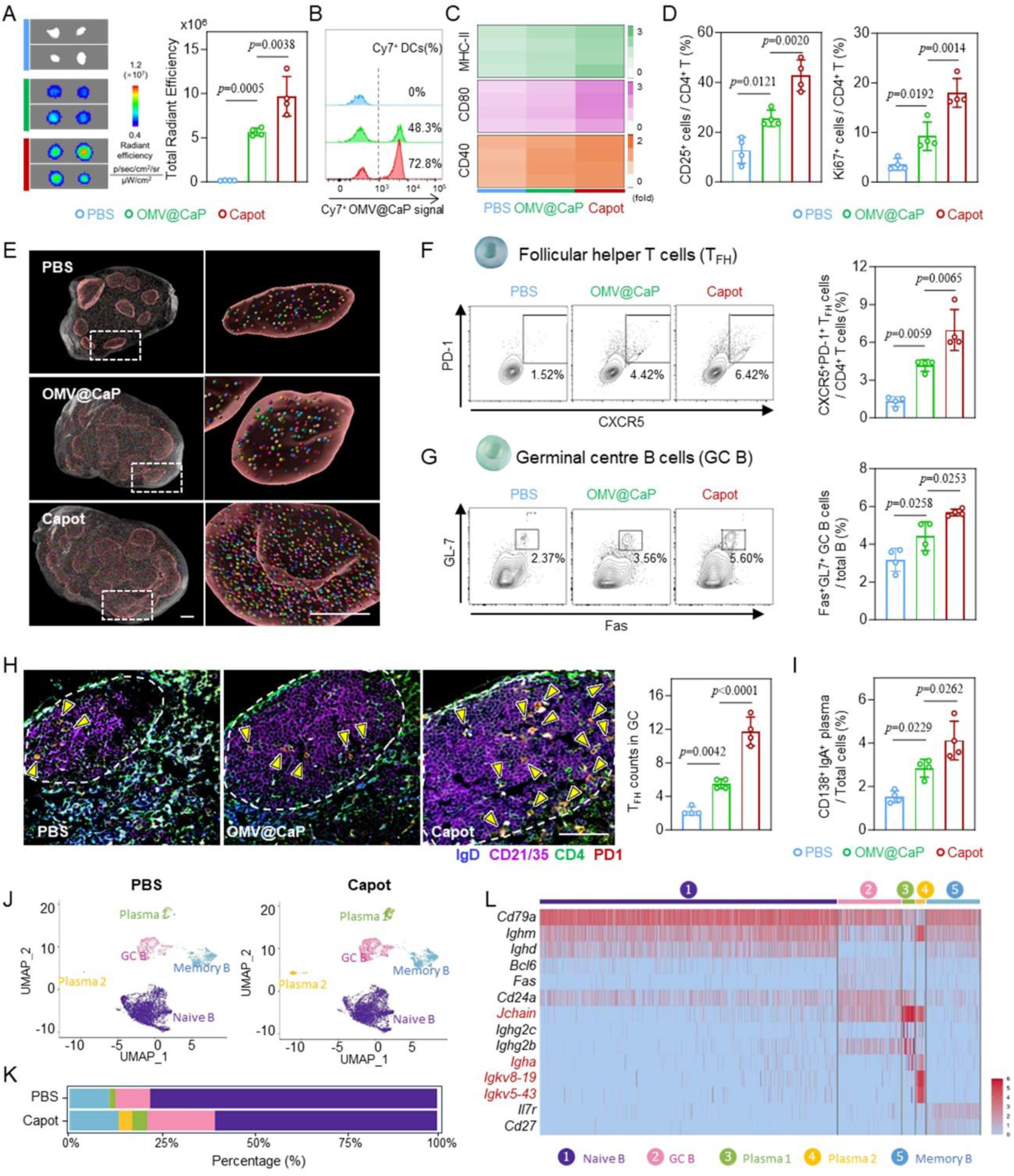
Immune activation of sLNs following sublingual vaccination in mice. A) Representative fluorescence images of sLNs at 24 h after sublingual administration of PBS, Cy7-labelled OMV@CaP or Capot (left), with corresponding quantitative fluorescence intensity (right; n = 4). B) Representative flow cytometry analysis showing the percentage of Cy7⁺ cells among CD11c^+^ DCs in sLNs. C) Expression levels of MHC-II, CD80, and CD40 on CD11c^+^ DCs in sLNs. D) Percentages of CD25^+^ (activation index, left) and Ki67^+^ (proliferation index, right) cells among CD4^+^ T cells in sLNs (n = 4). E) Three-dimensional images of light-sheet microscopy data showing B cell follicles (big red ball) and CD4^+^ T cells (small colorful dots) infiltration in sLNs. Scale bar = 100 μm. F) Representative flow cytometry plots (left) and quantification (right; n = 4) of follicular helper T cells (T_FH_, CXCR5^+^PD-1^+^) among CD4^+^ T cells. G) Representative flow cytometry plots (left) and quantification (right; n = 4) of germinal center B cells (GC B, Fas^+^GL7^+^) among total B cells. H) Representative sLN images after multiplex immunofluorescent staining for IgD, CD21/35, CD4, PD-1 (left), and corresponding quantification of CD4^+^PD1^+^ T_FH_ counts within CD21/35^+^IgD^-^ GC regions (right; n = 4). Scale bar = 50 μm. I) Percentages of CD138^+^IgA^+^ plasma cells in sLNs (n = 4). J) Uniform manifold approximation and projection (UMAP) plots of single-cell RNA sequencing data showing annotated B-cell clusters. K) Proportions of B-cell clusters identified in (J). L) Heatmap of the indicated genes across B-cell clusters identified in (J). Data in (A, D, F-I) are presented as means ± SD. Statistical significance was tested using one-way ANOVA with multiple comparison tests (A, D, F-I). *p* values have been presented in the figure.

We next evaluated T follicular helper (T_FH_) cells, which provide specialized help to B cells and are indispensable for germinal centre (GC) formation, affinity maturation, and the production of high-affinity antibodies^32^. Flow cytometry revealed that both OMV@CaP and Capot immunization markedly increased the proportion of T_FH_ cells (CD3⁺CD4⁺CXCR5⁺PD-1⁺) among CD4 T cells (Figure 2F), accompanied by a corresponding rise in the frequency of GC B cells (B220⁺CD3⁻Fas⁺GL7⁺) among total B cells (Figure 2G). To further visualize T_FH_ infiltration into the GC, sLNs from the three groups were paraffin-embedded and stained with fluorescent antibodies. Compared with the PBS group, both OMV@CaP and Capot increased T_FH_ (CD4⁺PD-1⁺) counts the GC region (CD21/35⁺IgD⁻) (Figure 2H). In addition, plasma cell differentiation was promoted, evidenced by a significant rise in IgA⁺ plasma cells (Figure 2I). Notably, across all these parameters, Capot consistently outperformed OMV@CaP, highlighting the advantage of the tablet-based formulation in enhancing bioavailability and amplifying coordinated immune responses within the sLNs.

To further dissect the B-cell response, we performed single-cell RNA sequencing of B cells from PBS- and Capot-immunized sLNs (Figure 2J, Figure S5A). The analysis confirmed an expanded GC B-cell population in the Capot group (Figure 2K), with these cells showing elevated expression of activation-associated genes such as *Egr1* and *Cited2* compared to PBS controls (Figure S5B), indicating a heightened activation state ^33,34^. Capot-derived plasma cells also displayed stronger activation, evidenced by increased expression of *Mzb1* (linked to antibody synthesis and secretion) and *Ccr10* (guiding plasma cell migration to mucosal and peripheral tissues) (Figure S5C) ^34,35^. Notably, a distinct plasma cell cluster (plasma 2) emerged exclusively in the Capot group, suggesting the generation of an OMV-specific IgA-producing plasma cell population, characterized by high expression of *J chain*, *Igha*, *Igkv8-19*, and *Igkv5-43* (Figure 2L). These transcriptional features suggest that Capot supports the emergence of specialized IgA-producing plasma cell states associated with mucosal immunity.

### Induction of oral mucosal antibody responses by Capot

The expansion of IgA⁺ plasma cell populations prompted us to directly quantify antibody responses in immunized mice. After optimization of the dosing regimen (Figure S6A), a two-dose schedule at a 2-week interval was selected to quantify *P.g* specific antibodies in saliva. At 14 days after the second immunization, Capot group showed obvious increase in salivary *P.g-*specific sIgA antibody and IgG antibody (Figure S6B). Given the dominant antibacterial role of salivary sIgA in the oral cavity (Figure 1B-E), we next performed longitudinal analyses of salivary sIgA, which revealed that Capot sustained high salivary sIgA titers for more than one year (Figure 3A).

**Figure 3.**
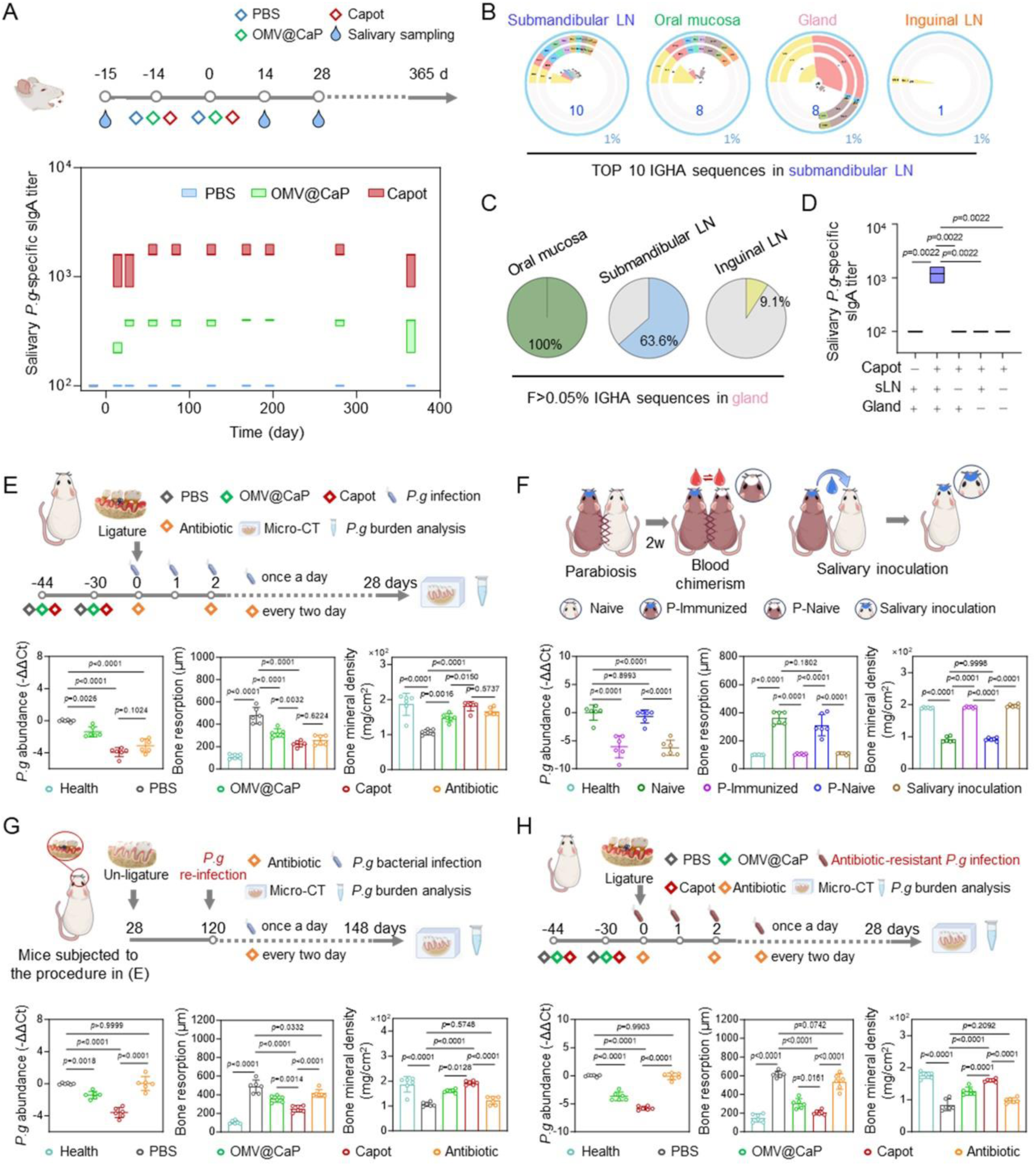
Mucosal antibody response of Capot and its protection efficacy across multiple mice periodontitis models. A) Schematic of the immunization and sampling schedule in mice (top) and longitudinal quantification of salivary *P.g*-specific sIgA titers at the indicated time points (bottom; n = 6). B) Clonal homology analysis of high-frequency BCR sequences (IGHA) in sLN compared with oral mucosa, salivary gland, and inguinal LN following Capot administration. C) Clonal homology analysis of high-frequency BCR sequences (IGHA) in salivary gland compared with oral mucosa, sLN, and inguinal LN following Capot administration. D) Salivary *P.g*-specific sIgA levels in Capot-immunized mice subjected to sham surgery, sLN excision, salivary gland excision, or combined excision, with unimmunized mice as control (n = 6). E) Schematic of immunization and primary periodontitis model establishment timeline (top), with quantification of salivary *P.g* burden by qPCR and micro-CT based analysis of alveolar bone loss on day 28 (bottom; n = 6). F) Schematic of parabiosis or salivary inoculation models (top), with corresponding salivary *P.g* burden and micro-CT analysis in the indicated models after immunization and primary periodontitis induction (bottom; n = 6). G) Schematic of secondary reinfection periodontitis model establishment timeline (top). Mice underwent the same procedure as in (E) before day 28, followed by reinfection. Salivary *P.g* burden and micro-CT analyses were performed on day 148 (bottom; n = 6). H) Schematic of immunization and antibiotic-resistant *P.g* induced periodontitis model establishment timeline (top), with salivary *P.g* burden and micro-CT analysis on day 28 (bottom; n = 6). Box-and-whisker plots in (A, D) show the median (central line), with the top and bottom of the box representing the maximum and minimum values. Data in (E-H) are presented as means ± SD. Statistical significance was tested using Kruskal-Wallis test with multiple comparison tests in (D) and one-way ANOVA with multiple comparison tests (E-H). *p* values have been presented in the figure.

After confirming that Capot induces IgA⁺ plasma cells in sLNs and promotes salivary sIgA, we examined how these two compartments are connected. Given that IgA⁺ plasma cells in sLNs express elevated *Ccr10* and are likely to migrate to peripheral tissues (Figure S5C), we analyzed BCR clonotypes in sLNs, salivary glands, and oral mucosa from Capot-immunized mice, with inguinal LNs as a distal control (Figure 3B). Clonal homology analysis of IGHA CDR3 sequences revealed minimal overlap between sLNs and inguinal LNs (one of the top 10 sLN clonotypes), whereas eight of the top 10 sLN clonotypes were detected among the top 1% high-frequency sequences in salivary glands and oral mucosa. Moreover, these shared clonotypes constituted the majority of dominant IGHA sequences in glands. Alternatively, using high-frequency gland IGHA sequences for cross-tissue comparison, we found clonal overlap rates of 100% with oral mucosa, 63.6% with sLNs, and only 9.1% with inguinal LNs (Figure 3C). To functionally assess the contribution of salivary glands in the salivary sIgA secretion, we selectively removed sLNs, salivary glands, or both prior to Capot immunization (Figure S7A). In all three settings, the induction of salivary sIgA was abolished (Figure 3D). Together with the high expression of the IgA transporter polymeric immunoglobulin receptor (pIgR) in the ductal epithelium of salivary glands (Figure S7B), these results identified the salivary gland as an essential site for detectable salivary sIgA secretion.

### Protective efficacy of Capot across multiple periodontitis models

We next examined whether Capot immunization could prevent *P.g*-induced periodontitis. Ligature-induced periodontitis model by placing silk ligatures around molars and daily inoculating them with *P.g* was established in mice pre-treated twice with PBS, OMV@CaP, or Capot (Figure 3E). As a positive control representing the current clinical standard of care, a local antibiotic treatment regimen was included, consisting of repeated administrations every two days for a total of 14 doses. After 28 days of infection, qPCR of salivary *P.g* abundance revealed marked bacterial reduction in the OMV@CaP, Capot, and antibiotic groups compared with PBS controls. Notably, *P.g* levels in the Capot and antibiotic groups were comparable. Consistently, micro-CT analysis demonstrated that *P.g* infection induced substantial alveolar bone resorption and bone mineral density loss, both of which were strongly suppressed in the Capot and antibiotic groups, yielding comparable therapeutic efficacy (Figure 3E and Figure S8). Histological and Tartrate-Resistant Acid Phosphatase (TRAP) staining analyses corroborated these findings (Figure S8).

To dissect whether mucosal or systemic antibodies mediated this protection, we generated four mouse cohorts with distinct antibody profiles *via* parabiosis or salivary inoculation between Capot-immunized (serum Ig⁺ / saliva Ig⁺) and naive (serum Ig⁻ / saliva Ig⁻) animals^36^. The resulting groups included naive (serum Ig⁻ / saliva Ig⁻), Parabiosis (P)-immunized (serum Ig⁺ / saliva Ig⁺), Parabiosis (P)-naive (serum Ig⁺ / saliva Ig⁻), and salivary-inoculation (serum Ig⁻ / saliva Ig⁺) mice (Figure 3F and S9A,B). Following *P.g* infection, qPCR and micro-CT results showed that P-naive mice exhibited bacterial burden and bone loss comparable to naive PBS controls (Figure 3F and Figure S9C), indicating that serum antibodies alone did not confer protection. In contrast, salivary-inoculated mice achieved periodontitis prevention comparable to P-immunized mice, demonstrating that mucosal rather than systemic antibodies mediated the protective effect, consistent with observations in clinical patients (Figure 1B-E).

Given that Capot immunization elicited durable salivary antibody responses lasting over one year (Figure 3A), we next examined whether this immunity could provide long-term protection against recurrent infection. To model disease relapse, ligatures were removed on day 28 after the initial infection and re-applied three months later, followed by *P.g* reinoculation (Figure 3G). Remarkably, qPCR, micro-CT, and histological analyses all demonstrated that Capot immunization provided robust protection against *P.g* reinfection and periodontitis recurrence (Figure 3G and Figure S10). In contrast, the antibiotic group showed no significant difference from PBS controls during reinfection, highlighting the superiority of Capot-induced long-term immune protection over transient antibiotic therapy.

Finally, considering the growing clinical challenge of antibiotic resistance, we evaluated Capot’s efficacy against *P.g* strain resistant to antibiotic (minocycline hydrochloride) (Figure 3H). The resistant strain displayed an approximately 75-fold increase in half-maximal inhibitory concentration (IC50) compared to the wild type (Figure S11A). In mice infected with this resistant *P.g*, antibiotic treatment failed to reduce bacterial burden or bone loss, whereas Capot immunization still provided strong protection against infection and periodontitis (Figure 3H, Figure S11B). These results underscore the potential of Capot as an effective immunization strategy capable of providing durable protection even against antibiotic-resistant bacteria.

### Immune responses and anti-periodontitis efficacy of Capot in non-human primates

To further assess the translational potential of Capot, we conducted immunization studies in cynomolgus monkeys (Figure 4A). Two doses of Capot were administered on day -14 and day 0, and the tablet completely disintegrated within one minute upon contact with saliva (Figure 4B). No local mucosal abnormity was observed after immunization, and the complete blood count and blood biochemical parameters remained within normal ranges (Figure 4C, Figure S12A,B). Photoacoustic imaging of the monkey treated with Cy7-labeled Capot revealed clear Cy7 signals inside the sLNs 24 hours after immunization (Figure 4D). Longitudinal ultrasound imaging further showed a gradually enlargement of the sLNs following Capot administration, returning to baseline size around 80 days (Figure 4E,F). Immunofluorescence staining of sLNs collected at day 28 demonstrated that, compared with PBS control, Capot immunization significantly increased the number of B cell zones and promoted greater CD4 T cell infiltration within these regions (Figure 4G,H), indicating robust activation of the sLN immune response. Antibody assays further showed that Capot induced obvious elevation in *P.g*-specific salivary sIgA antibody 14 days after the booster immunization (Figure 4I). Over time, salivary sIgA levels continued to rise during the first 100 days and remained stably for at least one year.

**Figure 4.**
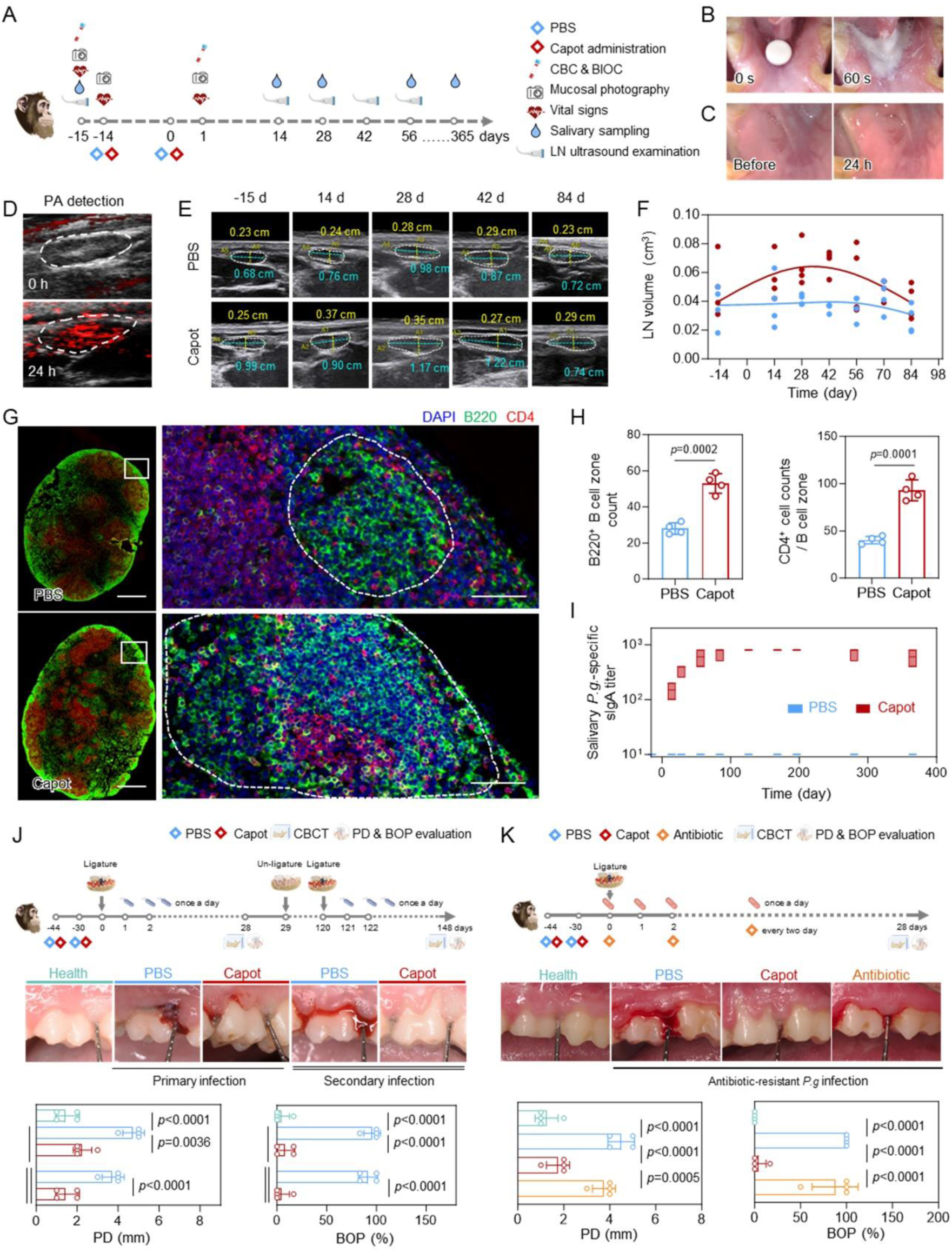
Performance of Capot in cynomolgus monkeys. A) Schematic illustration of the immunization regimen and sampling schedule in cynomolgus monkeys. CBC = complete blood count; BIOC = blood biochemistry. B) Representative photographs showing Capot disintegration in the oral cavity. C) Representative photographs of the sublingual mucosa before and 24 h after sublingual administration of Capot. D) Representative photoacoustic (PA) images of sLN before and 24 h after sublingual administration of Cy7-labelled Capot. E) Representative ultrasound images of sLN at the indicated time points following Capot administration. F) Quantitative analysis of sLN volume over time (n = 4). G) Representative sLN images following immunofluorescent staining for B220 (green), CD4 (red), and DAPI (blue). Scale bar = 1 mm (left) and 50 μm (right). H) Quantification of B220^+^ B cell zones and CD4^+^ T cell counts inside for (G) (n = 4). I) Salivary *P.g*-specific sIgA titers at the indicated time points (n = 4). J) Schematic of immunization, primary & secondary periodontitis models establishment (top), representative buccal tooth photographs (middle), and quantitative PD & BOP evaluation at day 28 (primary infection) and day 148 (secondary infection) (bottom; n = 4). K) Schematic of immunization and antibiotic-resistant *P.g*-induced primary periodontitis model establishment (top), representative buccal tooth photographs (middle), and quantitative PD & BOP evaluation at day 28 (bottom; n = 4). Data in (H, J, K) are presented as means ± SD. Box-and-whisker plots in (I) show the median (central line), with the top and bottom of the box representing the maximum and minimum values. Statistical significance was tested using unpaired two-tailed t-test (H) and one-way ANOVA with multiple comparison tests (J, K). *p* values have been presented in the figure.

To assess therapeutic efficacy, *P.g*-induced primary and reinfection periodontitis models were established in cynomolgus monkeys (Figure 4J). qPCR analyses of salivary samples at day 28 revealed markedly reduced *P.g* abundance in Capot-immunized monkeys compared with PBS controls (Figure S12C). Oral examination at day 28 showed that Capot effectively mitigated *P.g*-induced clinical deterioration, including reduced PD and BOP (Figure 4J), which accompanied by the reduced salivary inflammatory factors (Figure S12D). *In vivo* cone-beam computed tomography (CBCT) further confirmed that Capot significantly inhibited alveolar bone resorption and bone mineral density loss (Figure S12E). Consistent results were obtained at day 148 in the *P.g* reinfection model, demonstrating Capot’s durable protection against recurrent infection (Figure 4J, Figure S12C-E). Finally, in a periodontitis model induced by antibiotic-resistant *P.g* infection, Capot immunization conferred robust protection against periodontal tissue destruction, whereas antibiotic treatment showed minimal efficacy, as assessed by oral examination and CBCT analyses (Figure 4K and S12F). Together, these results establish Capot as a safe and potent mucosal vaccine capable of eliciting long-lasting immune responses and conferring robust protection against primary, recurrent, and antibiotic-resistant periodontitis in nonhuman primates.

## Discussion

This study establishes a sublingual tablet vaccine as a practical strategy for activating oral mucosal immunity. Guided by clinical observations linking higher salivary *P.g*-specific sIgA levels with reduced bacterial burden and improved periodontal outcomes, we developed Capot, a rapidly disintegrating sublingual vaccine designed to elicit functional mucosal antibody responses. Across mice and non-human primates, Capot consistently elevated salivary sIgA and enhanced saliva-mediated antibacterial activity, with protective effects demonstrated in experimental models of primary, recurrent, and antibiotic-resistant oral infection.

The design of Capot integrates bacterial extracellular vesicles with a CaP coating and an orally disintegrating tablet format to coordinate immunogenicity, safety, and delivery efficiency. Bacterial vesicles intrinsically contain a broad repertoire of native antigens and multiple PAMPs, enabling coordinated immune activation without predefined antigen selection or exogenous adjuvants. In our experimental setting, however, unmodified vesicles elicited local mucosal irritation, indicating a limitation for their direct application at oral surfaces. Encapsulation within a CaP shell mitigates this liability by providing a biocompatible interface that dampens local inflammatory responses while preserving immunogenic content. Formulation as an orally disintegrating tablet further improves delivery efficiency by enabling rapid disintegration in saliva, limiting gastrointestinal exposure, and enhancing local availability at the oral mucosa. Compared with injected whole-cell vaccines, Capot enables non-invasive immunization while directing immune activation toward the sLNs, thereby preferentially eliciting oral mucosal antibody responses. Relative to multivalent subunit vaccines that typically combine selected antigens with exogenous adjuvants^25^, vesicle-based immunization preserves native antigenic and adjuvant complexity in a single nanoscale formulation, reducing manufacturing complexity and the risk of epitope omission. Unlike oral or intranasal antibacterial vaccines that primarily engage gastrointestinal or respiratory mucosa^19,25^, Capot is designed to preferentially target the oral mucosal immune compartment, providing a focused strategy for protection against oral infectious diseases like periodontitis.

The translational potential of Capot is supported by several features relevant to clinical deployment. The materials used—including lyophilized vesicles, CaP and standard excipients commonly employed in sublingual orally disintegrating tablets—exhibited favorable safety profiles across two tested species, with no evidence of overt mucosal damage or systemic toxicity. Capot also demonstrated excellent physicochemical stability following freeze-drying, retaining immunogenic activity after extended storage at room temperature. This thermostability reduces reliance on cold-chain logistics and is particularly advantageous for vaccine distribution in regions with limited refrigeration infrastructure. Importantly, the consistent induction of bacteria-specific salivary sIgA responses observed across mice and non-human primates highlights the oral mucosal immune compartment as a conserved and accessible target for vaccination. Combined with its needle-free, self-administrable format, Capot represents a scalable and compliance-friendly strategy for public health applications.

From a clinical perspective, Capot addresses a preventive need distinct from that of antibiotic therapy. While antibiotics remain essential for treating acute infections, their effectiveness against chronic mucosal disease is increasingly constrained by recurrent exposure and the global rise of drug-resistant bacteria^37^. Vaccination strategies that enhance mucosal immune surveillance can reduce bacterial burden and suppress pathogenic activity before disease exacerbation. In this context, Capot is well suited for preventing or mitigating chronic oral infections through sustained local antibody responses, without relying on direct antimicrobial mechanisms.

## Methods

### Data reporting

No statistical methods were used to predetermine sample size. Histopathology slides were read by veterinary pathologists blinded to treatment groups; all other studies were performed unblinded as analysis results could not be manipulated by interpretation.

### Animals

C57BL/6 mice (6-8 weeks old) were procured from Beijing Vital River Laboratories and randomly allocated into distinct experimental groups for subsequent assays. The experimental protocol was reviewed and approved by the Animal Ethics Committee of the Institute of Process Engineering (approval ID: IPEAECA2021083).

Cynomolgus monkeys (4-5-years old) were obtained and housed at the Animal Center of the Academy of Military Medical Sciences. All related experiments were approved by the Institutional Animal Care and Use Committee of the Academy of Military Medical Sciences (Approval ID: IACUC-DWZX-2022-057) and conducted in line with its institutional guidelines.

All animal procedures in this study were conducted in strict compliance with the Regulations for the Care and Use of Laboratory Animals and the National Standard for Ethical Review of Animal Experiments (GB/T 35892-2018, China).

### Cell lines and primary cells

*Porphyromonas gingivalis* (*P.g*, ATCC33277, purchased from Rayzbio, Shanghai) was cultured on brain heart infusion (BHI) agar medium containing sterile defibrinated sheep blood and hemin-vitamin K (Solarbio, Beijing, China), and anaerobically incubated at 37°C for 7-10 days until colonies with a standard morphology are clearly visible. Single colonies were picked and inoculated into brain heart infusion liquid medium containing hemin-vitamin K (Solarbio), followed by anaerobic culture at 37°C until the logarithmic growth phase (2.5×10^11^ CFU per mL).

Bone marrow derived dendritic cells (DCs) were generated from the bone mesenchymal stem cells (BMSCs) of C57BL/6 mice by induced differentiation. Briefly, the BMSCs were flushed from mouse marrow cavities of femurs and tibias and then cultured in the RPMI 1640 medium containing 10% FBS and 1% penicillin/streptomycin in the presence of recombinant mouse granulocyte- macrophage colony-stimulating factor (GM-CSF) (20 ng/ml) and IL-4 (10 ng/ml). After 5 days, BMDCs were harvested for further use.

### Human samples

Saliva and serum samples from clinical periodontitis patients were provided by Beijing Stomatological Hospital, Capital Medical University. Written informed consent was obtained from each individual for sample collection. This study was reviewed and approved by the Clinical Ethics Committee of Beijing Stomatological Hospital, Capital Medical University (Approval ID: CMUSH-IR-KJ-PJ-2023-15).

### Fabrication and characterization of the Capot

#### Preparation of OMV

The bacterial culture medium was subjected to centrifugation at 8,000 g for 20 minutes to eliminate bacterial cells, after which it was filtered *via* a 0.22 μm poresize filter. Subsequently, the medium was centrifuged at 110,000 g for 3 hours at 4 °C (XPN, Beckman Coulter). The OMV pellet obtained was resuspended in 0.90% (w/v) NaCl and filtered through a 0.22 μm poresize filter to prevent contamination by bacteria or cell debris. The particle counts of OMV was determined using the Exoid (TRPS) nanoparticle bioanalyzer (Izon Science, Oxford, UK). Samples were stored at -80 °C for subsequent experiments.

#### Preparation of OMV@CaP

OMVs (1.5 × 10^12^ particles) were dispersed in 1 mL of DMEM medium (Gibco) and placed on a vertical mixer at 4 °C for overnight incubation to achieve equilibrium. Next, 10 µL of CaCl₂ (1 M) was added to the reaction system, which was then incubated at 37 °C for an additional 2 hours. Following incubation, the biomineralized OMVs (designated as OMV@CaP) were rinsed three times with ultrapure (UP) water via centrifugation at 14,000 g for 10 minutes each time. The resulting pellet was resuspended in NS solution or UP water for subsequent characterization.

#### Characterizations of OMV and OMV@CaP

For TEM observations, OMV sample was initially fixed in 1% glutaraldehyde for 15 minutes. Subsequently, the OMV sample was deposited onto 200-mesh copper grids and stained with 2% uranyl acetate for 30 seconds, after which they were rinsed twice with distilled water. Imaging was performed using a JEM-1400 Flash microscope (JEOL) operated at 120 kV. For analyses of size and zeta potential, OMV and OMV@CaP sample was examined *via* dynamic light scattering (DLS; Nanoseries, Malvern).

In terms of TEM imaging, a suspension of OMV@CaP was dropped onto a carbon film-supported copper grid and allowed to dry at room temperature. The structure of OMV@CaP was directly observed without any staining using a JEM-1400 Flash microscope (JEOL) operated at 120 kV. The element mapping analysis was performed using FEI Talos F200X operated at 200 kV. For thermogravimetric and differential scanning calorimetry (TG-DSC) analysis, OMV@CaP sample was lyophilized and then measured using an SDT-Q600 instrument. The analysis was conducted at a heating rate of 10 °C per minute, ranging from 25 °C to 800 °C under a nitrogen atmosphere. To assess the *in vitro* release of Ca²⁺, OMV@CaPs were suspended in normal saline solutions with different pH values (5.0 and 7.0). Supernatants were collected at different time points *via* centrifugation and Ca²⁺ was detected using an MP523-03 calcium ion counter.

#### Preparation of Capot

Lyophilized OMV@CaPs was compressed into tablets together with pharmaceutical excipients (62% microcrystalline cellulose, 20% croscarmellose sodium, 16% starch, 2% micronized silica gel) using the appropriate mold.

### Safety evaluation

#### Visualization of vesicle trafficking

For mice, OMVs were labeled with Cy7 and further formulated into Cy7-labeled OMV@CaP and Capot. Above three Cy7-labeled vaccines with equivalent total particle counts were separately administered sublingually in different mice. The digestive tract tissues of mice were isolated 24 h after immunization, and the submaxillary lymph nodes of mice were harvested at 12, 24, 48 and 72 h after immunization, which were imaged using IVIS Spectrum Imaging System (PerkinElmer). For cynomolgus monkeys, photoacoustic detections of sLN were performed before and 24 h after sublingual administration of Cy7-labelled Capot (LAZR-X Vevo, FUJIFILM VisualSonics, Canada).

#### Inflammation evaluation

At 24 h after sublingual immunization, the occurrence of redness and swelling was observed under the stereoscopic microscope (Nikon SMZ800N). Local histological changes were evaluated using hematoxylin and eosin (H&E) staining. The expression levels of TNF-α, IL-1β, IL-6, and IFN-γ were determined using ELISA.

#### Visualization of *In Vivo* IFN-γ Production

At various time points, IFN-γ-IRES Venus-AkaLuc mice that received sublingual administration of OMVs, OMV@CaPs, or Capot were intraperitoneally injected with 100 µL of 5 mM Aka Lumine n-Hydrochloride (Wako) 15 minutes prior to imaging. Imaging was performed using the IVIS Spectrum Imaging System (PerkinElmer).

### Immunofluorescence analysis

#### Light-sheet imaging

For visualizing the CD4⁺ T cell and B cell location in sLNs, CD4^+^ T cells and B cells were stained with anti-CD4-FITC (GK1.5, Biolegend, 100406) and anti-B220-AF647 (RA3-6B2, Biolegend, 103229), respectively. The sLNs were processed with tissue-clearing reagent and captured in CUBIC-R+ reagent by light-sheet microscopy (Light Innovation Technology). Laser power and gain were adjusted consistently for each group. Image reconstructions were generated using LitScan v3.0 software.

For three-dimensional (3D) imaging data acquirements, segmentations of target structures, including sLNs, B220^+^ B cell zones and CD4^+^ T cells were performed using the Interactive Thresholding module and the segmentation workroom in Amira software (Thermo Fisher Scientific). The 3D imaging analysis results are rendered and displayed using the Imaris software (Oxford Instruments).

#### Immunofluorescence analysis of tissue sections

Mice sLNs were fixed with 4% paraformaldehyde for 12 hours at 4 ℃ and dehydrated in 30% sucrose overnight before being embedded in OCT (Sakura). OCT-embedded 20 μm cryostat sections were blocked in PBS with 0.3% Triton and 5% FBS before staining. Slides were then stained with anti-IgD-ef450 (Invitrogen, 48-5993-82, 1:150), anti-CD21/35-PE (BD, clone 7G6, 1:150), anti-CD4-FITC (Biolegend, 100406, 1:150), and anti-PD1-APC (biolegend, 109111, 1:150) antibodies. Tissue sections were mounted with the ProlongGold Antifade reagent (Invitrogen).

Cynomolgus monkey sLNs were fixed in 4% paraformaldehyde solution, followed by paraffin embedding and subsequent serial sectioning. 5 µm thick sLN sections were prepared for immunofluorescence staining assay. Antigen retrieval was performed via microwave treatment with ethylenediaminetetraacetic acid (EDTA) retrieval buffer, and then the sections were blocked with 5% (w/v) bovine serum albumin for 30 minutes. The sections were incubated with anti-B220 antibody (Abcam, ab59168, 1:150) and anti-CD4 antibody (Invitrogen, MA5-47842, 1:150) at 4 °C overnight. Thereafter, the corresponding fluorescent secondary antibodies (Abcam, ab150115, 1:10,000; Abcam, ab50168, 1:10,000) were added for 1-hour incubation at room temperature. DAPI was used for nuclear counterstaining, and the stained sections were mounted and imaged on the Vectra imaging platform.

### Flow Cytometry Analysis

#### Analysis of vaccine uptake by DCs in sLNs

After administration with Cy7-labeled OMVs, OMV@CaP, or Capot, single-cell suspensions from the sLNs were prepared, following centrifugation and resuspension with staining buffer. The collected cells were stained with anti-CD11c-AF700 (N418, Biolegend, 117320, 1:150) to define dendritic cells. The proportion of DCs that internalized particles (Cy7^+^CD11c^+^) was measured using CytoFLEX LX flow cytometer (Beckman Coulter), and analyzed using CytExpert software (v. 2.3).

#### Analysis of DCs activation in sLNs

Single-cell suspensions from the sLNs were prepared, following centrifugation and resuspension with staining buffer. The collected cells were stained with anti-CD11c-BV650 (N418, Biolegend, 117339, 1:150) and Zombie Yellow™ Fixable Viability Kit (Biolegend, 423103) to define live DCs, and anti-CD40-PE (clone FGK45, Biolegend, 157506, 1:150), anti-CD80-APC (clone 16-10A1, Biolegend, 104713, 1:150), and anti-MHC II-Alexa Fluor 700 (clone M5/114.15.2, Biolegend, 107622, 1:150) as activation markers. The mean fluorescence intensity was measured using CytoFLEX LX flow cytometer (Beckman Coulter), and analysed using CytExpert software (v. 2.3).

#### Analysis of T cell activation in sLNs

Single-cell suspensions from the sLNs were prepared, following centrifugation and resuspension with staining buffer. The collected cells were stained with anti-CD4-FITC (clone GK1.5, Biolegend, 100406, 1:150) and Zombie Yellow™ Fixable Viability Kit (Biolegend, 423103) to define live CD4^+^ T cells, and anti-CD25-PE (clone A18246A, Biolegend, 113704, 1:150), anti-Ki67-APC (clone 16A8, Biolegend, 652406, 1:150) as activation markers.

In addition, the proportion of T_FH_ cells was detected by staining with anti-CD4-FITC (clone GK1.5, Biolegend, 100406, 1:150), Zombie Yellow™ Fixable Viability Kit (Biolegend, 423103), anti-CXCR5-Brilliant Violet 421 (clone L138D7, Biolegend, 145511, 1:150), anti-PD-1-PE (clone 29F.1A12, Biolegend, 135205, 1:150).

#### Analysis of GC B cell and IgA^+^ plasma cell in sLNs

Single-cell suspensions from the sLNs were prepared, following centrifugation and resuspension with staining buffer. The proportion of GC B cells was detected by staining with anti-B220-FITC (clone RA3-6B2, Biolegend, 103205, 1:150), Zombie Yellow™ Fixable Viability Kit (Biolegend, 423103), anti-Fas-APC (clone SA367H8, Biolegend, 152603, 1:150), anti-GL7-PE (clone GL7, Biolegend, 144607, 1:150).

The proportion of IgA^+^ plasma cell was detected by staining with Zombie Yellow™ Fixable Viability Kit (Biolegend, 423103), anti-CD138-PE (clone W20051E, Biolegend, 112503, 1:150), and anti-IgA-FITC (clone mA-6E1, Invitrogen, 11-4204-83, 1:150).

### Single-cell RNA sequencing and BCR Repertoire Sequencing

#### Single-cell RNA sequencing

The single-cell suspensions were assessed for cell viability before proceeding with library construction using the Chromium Next GEM Single Cell 3′ Kit v3.1 (10x Genomics) according to the manufacturer’s instructions. Sequencing was performed on the Illumina NovaSeq 6000 platform. Raw BCL data were converted to FASTQ files and processed using the Cell Ranger pipeline (v.3.0) to generate gene-barcode matrices. Downstream analysis was performed in R (v.4.4.0) using Seurat (v.4.3.0). High-quality cells were retained by applying stringent filtering criteria (nFeature_RNA between 200 and 5000, mitochondrial content < 15%). The data were normalized (LogNormalize), scaled, and integrated using Harmony (v.1.2.1) to correct for batch effects. Principal component analysis (PCA) was performed, and the resulting embeddings were integrated using Harmony (v.1.2.1) to correct for batch effects. Dimensionality reduction was performed using UMAP, and clusters were identified via the Louvain algorithm (resolution = 0.5).

High-resolution characterization of the B cell population facilitated the annotation of specific types, including Naive B cells, Germinal Centre B cells, Memory B cells, and two populations of Plasma cells. The expression of marker genes for B cell types was visualized using a heatmap with Z-score scaled expression values. The proportional distribution of the annotated cell types between PBS and Capot groups was visualized using stacked bar charts.

#### BCR Repertoire Sequencing

RNA samples were isolated from superficial inguinal lymph nodes (sLNs), inguinal lymph nodes, oral tissues and glandular tissues after immunization, and subjected to in-depth analysis *via* high-throughput immunoglobulin heavy chain (IGH) sequencing using the ImmuHub BCR profiling system (ImmuQuad Biotech). Briefly, a 5′-RACE (rapid amplification of cDNA ends) protocol was employed to ensure unbiased amplification. During cDNA synthesis, unique molecular barcodes (UMBs) were incorporated to mitigate amplification bottlenecks and eliminate PCR/sequencing artifacts. Sequencing was performed on an Illumina NovaSeq platform in PE150 mode. A universal adapter containing UMBs was ligated to the 5′ end of first-strand cDNA, while reverse primers targeting IGH constant regions were designed to enable less biased PCR amplification. Raw sequence reads were processed using UMBs to correct errors and remove PCR duplicates. V, D, J, and C gene segments were annotated using IMGT, and CDR3 regions were extracted for clonotype assembly. Out-of-frame sequences and those containing premature stop codons were filtered from the IGH repertoire. Clonotype abundance was determined by aggregating counts of IGH clones sharing identical CDR3 nucleotide sequences. Clonal homology analysis was performed by calculating the percentage of perfectly matched paired sequences in BCR sequencing data from different anatomical sites, including the V segment, D segment, J segment, C segment, CDR3 gene sequence, and CDR3 amino acid sequence.

### Immunization

#### Immunization for mice

C57BL/6 mice (6-8 weeks old) were immunized at day -14 and day 0 with PBS, OMV@CaP or Capot containing the equivalent of bacterial OMVs. Each group contained six mice. Saliva sample collection was performed on day -15, 14, 28, 56, 84, 126, 168, 196, 280 and 365, for the subsequent antibody detection.

#### Immunization for cynomolgus monkeys

A total of 8 cynomolgus monkeys (4-5-years old) were divided into PBS group and Capot group. Each group contained four animals. Cynomolgus monkeys in Capot group were immunized twice with Capot containing the equivalent of bacterial OMVs at day -14 and day 0. Ultrasonographic examinations of sLNs were performed on day -15, 14, 28, 42, 56, 70, and 84 using Ultrasound Diagnostic Instrument V70 (Shantou Institute of Ultrasonic Instruments Co., Ltd., Guangdong, China). Saliva sample collection was performed on day -15, 14, 28, 56, 84, 126, 168, 196, 280 and 365, for the subsequent antibody detection.

### sLN and salivary gland excision models establishment

After routine disinfection and anesthesia, the skin incision was performed and the flap was opened along the inferior border of the mouse mandible to fully expose the mandibular salivary glands, as well as the surrounding lymph nodes. For sLN excision, all lymph node tissues were identified and carefully excised, with the salivary glands preserved intact. For gland excision, only the glands were excised, with lymph node tissues preserved intact. For combined excision, both the lymph node tissues and the glands were excised. For control, sham operation was performed. Briefly, skin incision and flap elevation were conducted, without excision of any tissues. After the operation, the flap was tightly sutured.

### Antibody detection

The binding affinities of serum and saliva from mice and monkeys to *P.g* were evaluated using ELISA. Corning 96-well plates were coated with 2 μg/ml of *P.g* ultrasonic extract protein in coating buffer and incubated overnight at 4 °C. Following incubation, plates were blocked with 5% bovine serum albumin in PBS for 2 hours at 37 °C. Serially diluted serum or saliva samples were added to the plates and incubated for 1 hour at 37 °C. Plates were then washed five times with 1× PBS containing 0.05% Tween-20 (PBST). Subsequently, 100 µL of horseradish peroxidase-conjugated secondary antibodies were added: goat anti-mouse IgG (Abcam, ab6789, 1:10,000), goat anti-mouse IgA (Abcam, ab97235, 1:5,000), goat anti-monkey IgG (Abcam, ab112767, 1:10,000), or goat anti-monkey IgA (4 A Biotech, ABIN458287, 1:5,000), respectively. Plates were incubated at 37 °C for 30 minutes and washed five more times with PBST. Colorimetric development was performed using 3,3′,5,5′-tetramethylbenzidine for 10 minutes at room temperature, and the reaction was terminated with 2 M H₂SO₄. Absorbance at 450 nm was measured using a microplate reader. The endpoint titer was defined as the highest reciprocal dilution of the sample yielding an absorbance value exceeding 2.1 times the OD₄₅₀ of the PBS control group.

### Periodontitis model establishment

#### Periodontitis primary-infection model establishment

Mice were immunized on day -44 and day -30, with OMV@CaP or Capot containing the equivalent of bacterial OMVs. On day 0, salivary *P.g*-specific sIgA across groups were detected by ELISA to confirm successful immunization. After that, following anesthesia, a 5-0 silk ligature (Johnson & Johnson, New Brunswick, NJ, USA) was placed around the upper left second molar and maintained in situ for 28 days. Concurrently, animals received daily oral application of *P.g* (10⁹ CFU/day) at the ligated site. During this period, mice in the antibiotic group received oral antibiotic intervention (minocycline hydrochloride gel) every two days. After that, all animals were euthanized. Alveolar bone resorption was assessed *via* micro-computed tomography (micro-CT; SIEMENS, Kyoto, Japan). Bone loss parameters were quantified using NIH ImageJ software. H&E stating was used for the observation of bone resorption. Tartrate-resistant acid phosphatase (TRAP) staining was used for the observation and evaluation of osteoclasts.

The immunization and modeling process for cynomolgus monkeys was similar to that for mice. Animals were immunized on day -44 and day -30 with Capot. On day 0, salivary *P.g*-specific sIgA across groups were detected by ELISA to confirm successful immunization. After routine anesthesia, a 4-0 silk ligature was placed around the maxillary first molar, followed by daily *P.g* (10⁹ CFU/day) inoculation at the ligated site. 28 days later, cone-beam computed tomography (CBCT) was used to detect the alveolar bone destruction of the modeled teeth, and periodontal specialist examinations were performed to evaluate the local inflammatory conditions of the modeled teeth. The concentration levels of salivary TNF-α, IL-1β, and IL-6 were determined using ELISA.

#### Periodontitis secondary-infection model establishment

After immunization and establishing the primary periodontitis infection model in mice and cynomolgus monkeys, ligatures were removed on day 28 after modeling to enable periodontal tissue healing. The animals were then maintained under standard housing conditions until day 120, after which ligature ligation was repeated on the same tooth following the identical protocol for the primary periodontitis model, so as to simulate the recurrence of periodontitis induced by secondary *P.g-*infection. On day 148, the severity of periodontitis was evaluated, with all assessment items consistent with those for periodontitis primary-infection models.

#### Antibiotic-resistant *P.g*-infection model establishment

The procedure for antibiotic-resistant *P.g*-infection model establishment is consistent with the periodontitis primary-infection model establishment procedure, except that the infecting bacteria are minocycline hydrochloride-resistant *P.g*. The construction process of the antibiotic-resistant *P.g* bacteria is as follows:

*P.g* was anaerobically cultured, serially diluted to ∼1×10⁶ CFU/mL. The minimum inhibitory concentration (MIC) of minocycline hydrochloride was determined using the broth microdilution adapted from CLSI M11 under anaerobic conditions. The MIC was defined as the lowest drug concentration completely inhibiting bacterial growth. For resistance induction, uninduced *P.g* was inoculated at a 1:100 volume ratio into fresh BHI medium containing minocycline hydrochloride at 0.01 μg/mL, which represented 1/10 of the initial MIC. The culture was incubated anaerobically and passaged every 48–72 hours. Bacterial tolerance to a given concentration was confirmed when the bacterial density reached 1×10⁹ CFU/mL and remained stable over five consecutive generations. Before increasing the drug concentration, the MIC of the induced strain was measured alongside that of the passaged parental strain (passaged equivalently without induction). This step tracked resistance development while controlling for passage effects. A strain was considered resistant when it maintained stable growth for five generations at a drug concentration ≥10 times the initial MIC. In parallel, MICs of the induced and parental strains were assessed by agar dilution at each stage. Genetic stability was verified by subculturing the resistant strain for five generations in drug-free medium. Finally, the strain was purified and stored long-term at −80°C or in liquid nitrogen.

For the antibiotic treatments in above mice primary-infection model, mice secondary-infection model, mice antibiotic-resistant *P.g*-infection model, and cynomolgus monkey antibiotic-resistant *P.g*-infection model, the periodontitis modeling procedure was consistent with other groups, except that 2% minocycline hydrochloride ointment was topically applied to the oral cavity once every two days during the modeling period.

### Quantitative real-time PCR

Salivary genomic DNA was extracted using the Rapid Bacterial Genomic DNA Isolation Kit (Sangon Biotech, Shanghai, China) and evaluated for purity and concentration using a NanoDrop spectrophotometer (Thermo Fisher Scientific, Waltham, USA). Quantitative PCR was performed on a system using the SYBR Fast Advanced Master Mix (Life Technologies, Carlsbad, CA, USA). The reaction mixture (20 µL) contained 10 µL of master mix, 0.4 µM of specific primers (Sangon Biotech, Shanghai) targeting the *P.g* 16S rRNA gene (Forward: 5’-CCCATTCTTTCCCGTTCTCTT-3’; Reverse: 5’-CAGCACATCCTTCTTTTCGATG-3’), primers targeting general bacteria (Forward: 5’-ACTCCTACGGGAGGCAGCAG-3’; Reverse: 5’-ATTACCGCGGCTGCTGG-3’), and 20 ng of template DNA. Cycling conditions consisted of an initial denaturation at 95 °C for 30 s, followed by 39 cycles of 95 °C for 5 s and 60 °C for 30 s, the entire procedure was carried out on an Archimed X4 fluorescence quantitative PCR instrument (ROCGENE, Beijing, China). A melt curve analysis was performed to verify amplicon specificity.

### Parabiosis model and salivary-inoculation model establishment

#### Parabiosis model establishment

Parabiosis was established as previously described ^[33]^: mice with matched genetic background, comparable weight and size were cohoused for at least 2 weeks to ensure harmonious cohabitation. Before the modeling, one of these two mice was immunized with Capot, and the serum/salivary antibody detection was performed to evaluate the success of immunization. After that, The mice were anesthetized and placed back-to-back with shaved surfaces upward, longitudinal skin incisions were made from 0.5 cm above the elbow to 0.5 cm below the knee, and the skin was separated from the fascia to create a 0.5 cm free margin; the left and right olecranons as well as knee joints of the two mice were sutured with non-absorbable 3-0 sutures, and the adjacent skin edges were closed with continuous absorbable 5-0 Vicryl suture from ventral to dorsal elbow to knee with tight closure around joints and no gaps. Two weeks later, serum/salivary antibody detection was performed on these two mice to evaluate the success of model establishment, followed by subsequent periodontitis modeling.

#### Salivary-inoculation model establishment

After immunization, mouse saliva samples were collected. The collected saliva samples were processed by centrifugation at 3,000 × g for 10 min at 4 °C to remove debris and cell contaminants. The supernatant was then filtered through a 0.22 μm sterile filter to ensure sterility. Subsequently, the saliva samples were evenly spread to the oral cavity of the salivary inoculation recipient mice, with a supplementary dose of 20 μL.

To investigate the inoculation frequency, the half-life of sIgA post-inoculation was measured. Briefly, the concentration of sIgA antibody in the saliva collected at fixed time intervals were detected using the aforementioned method. Then, the half-life of sIgA in the oral cavity of mice was calculated. Referring to the experimental results, saliva was inoculated every 4 hours in the subsequent modeling process.

### Statistical Analysis

GraphPad Prism 9.0.0 and Origin 2023 software were employed for data visualization and statistical testing. Results are presented as mean ± standard deviation (SD).

For data meeting assumptions of normality and equal variance, significance was determined using two-tailed unpaired t-tests or one-way ANOVA with either Dunnett’s or Tukey’s multiple comparison tests. For antibody titer measurements, statistical comparisons were conducted via Mann-Whitney U tests or Kruskal-Wallis tests.

Correlation analyses utilized Spearman’s rank correlation coefficient, with results reported as *p* values and R² values.

Importance analysis of *P.g* specific antibody classes in periodontitis prognosis based on SHAP interpretation of a trained XGBoost model.

For Single-cell RNA sequencing analysis, split violin plots were utilized, and differential expression analysis was performed using the Wilcoxon rank-sum test to compare the expression abundance of specific genes between groups.

All experiments were repeated with at least three biologically independent in the manuscript, except for the modeling in cynomolgus monkeys.

## Data availability

All data supporting the findings of this study are available in the Article and its Supplementary Information and are also available from the corresponding author on request.

## Declaration of competing interest

The authors declare that they have no known competing financial interests or personal relationships that could have influenced the work reported in this study.

## Contributions

W. Wei, G.M., S.W., J.X., and Yi Liu conceived and designed the study; Yit. Liu, Y.D., X.Y., and Q.Z. collected and processed the clinical samples; Yit. Liu performed the *in vitro* and *in vivo* experiments; X.H. performed AI-based clinical data analysis; Yit. Liu and Q.C. conducted bacteria-related experiments; Yit. Liu, Q.C., and X.L. performed immunological assessments; Yit. Liu performed single-cell RNA sequencing and bioinformatic analyses; P.G., W. Wang, and Dongf. Wang provided suggestions for single-cell RNA sequencing. C.P., Y.Q., Dongs. Wang, and W.L. provided guidance and methodological support for the non-human primate disease modeling; W. Wei, Q.C., X.H., X.L., P.G., Y.Z., H.L., W. Wang, D.Z., L.G., Dongf. Wang, and S.W., discussed the results and gave advice; Yit. Liu, S.W., and W. Wei wrote the manuscript; W. Wei, G.M., S.W., J.X., and Yi Liu further revised the manuscript.

## Funding

This work was supported by the National Natural Science Foundation of China (T2225021 to W. Wei, T2394501 to G.M., 82372130 to S.W., 82571088 to Yi Liu, 82122015 to J.X., 82571094 to J.X., 82201053 to Yit. Liu), National Key Research and Development Program of China (2023YFC2307700 to W. Wei), CAS Project for Young Scientists in Basic Research (YSBR-083 to W. Wei), CAST Youth Talent Support Program (2024QNRC001 to S.W.), Strategic Priority Research Program of the Chinese Academy of Sciences (XDC0290303 to S.W.), Natural Science Foundation of Beijing, China (L2510103 to Yit. Liu), Beijing Physician Scientist Training Project (BJPSTP-2025-29 to Yit. Liu), the Basic-Clinical Joint Research and Cultivation Platform of Capital Medical University (JLPYPT2025012 to Yit. Liu), the Innovation Foundation of Beijing Stomatological Hospital, Capital Medical University (CXJJ25101 to Yit. Liu), the National Science and Technology Major Project (2025ZD1804202 to D.Z.).

## Supporting Information

**Figure S1.**
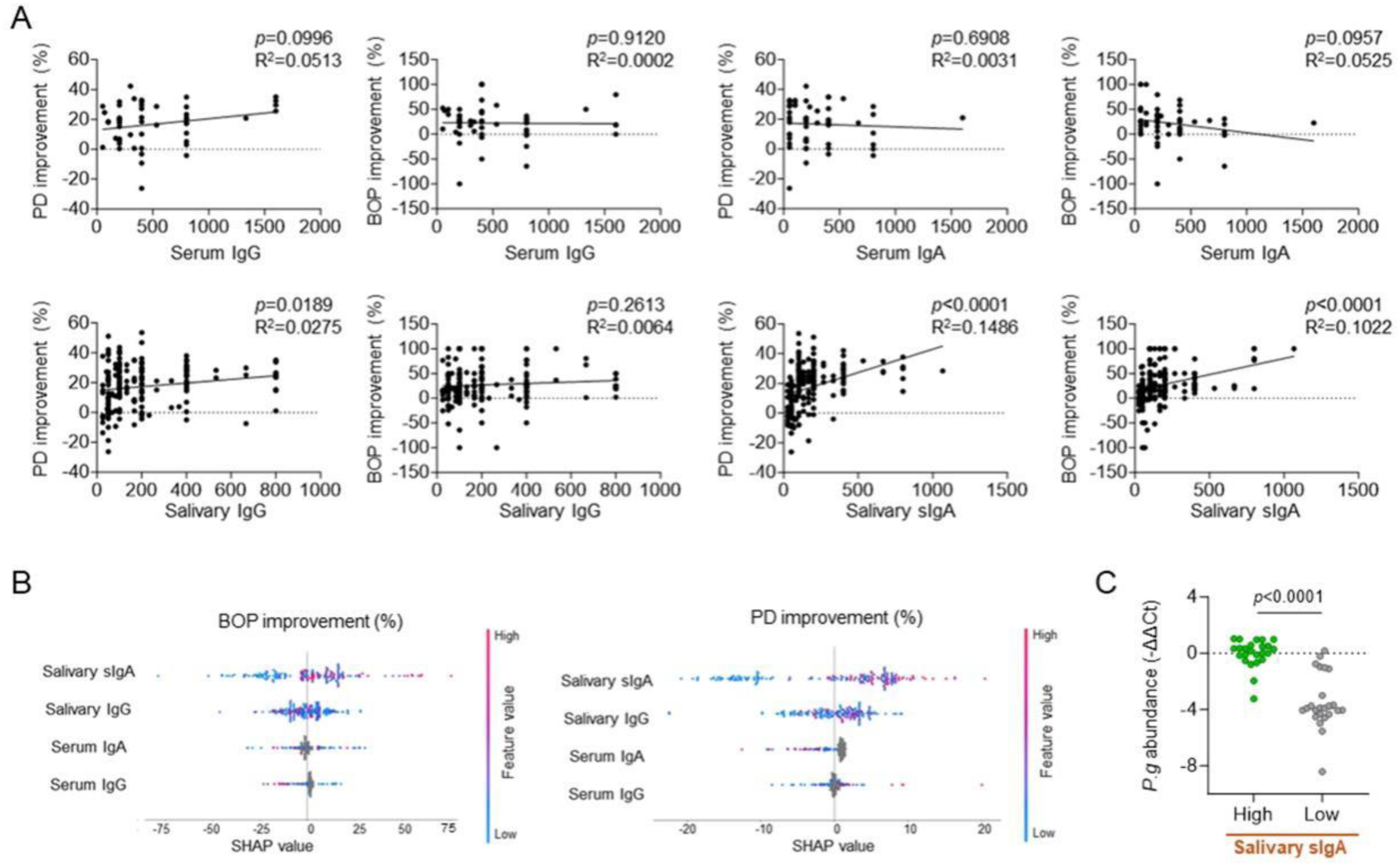
Clinical analysis for evaluating the association between antibody classes and periodontal outcomes. A) Pearson correlation analyses examining relationships between various antibody classes and periodontal outcomes, including serum *P.g*-specific IgG antibody titers (n = 54), serum *P.g*-specific IgA antibody titers (n = 54), salivary *P.g*-specific IgG antibody titers (n = 200), and salivary *P.g*-specific sIgA antibody titers (n = 200). B) SHAP (SHapley Additive exPlanations) interpretation of the antibody classes associated with periodontal outcomes based on XGBoost model. C) Quantitative PCR analysis of *P.g* burden in salivary from representative patients related to Figure 1E (n = 24). Statistical significance was tested with Pearson correlation analysis (A) and unpaired two-tailed t-tests (C). *p* values have been presented in the figure.

**Figure S2.**
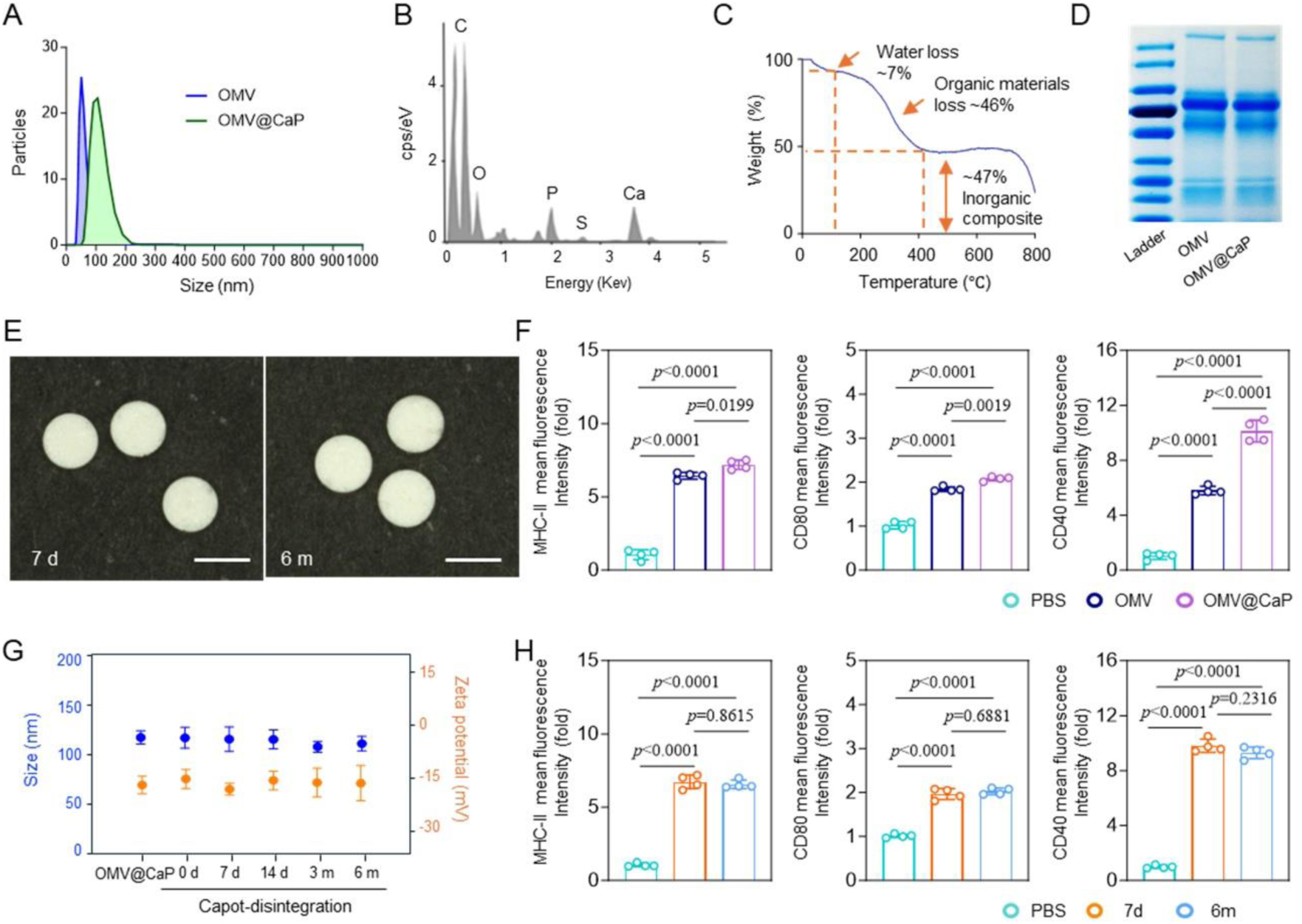
Preparation and characterization of OMV@CaP and Capot. A) Particle size analysis of OMV and OMV@CaP. B) Elemental analysis of OMV@CaP based on energy dispersive X-ray detector. C) Thermogravimetric and differential scanning calorimetry analysis of OMV@CaP. D) SDS-PAGE analysis of the protein in OMV and OMV@CaP. E) Representative photographs of Capot, which were stored at room temperature for 7 days (left) and 6 months (right). Scale bar = 2 mm. F) Flow cytometry analysis showing MHC-II, CD80, and CD40 expression levels on CD11c^+^ DCs in sLNs after sublingual administration of PBS, OMV or OMV@CaP (n = 4). G) Particle size and zeta potential of fresh prepared OMV@CaP and OMV@CaP released from different Capot formulations that storage at room temperature for 7 days, 14 days, 3 months, and 6 months (n = 4). H) Flow cytometry analysis showing MHC-II, CD80, and CD40 expression levels on CD11c^+^ DCs after exposure to released OMV@CaP in (G) (n = 4). Data in (F, G, H) are presented as means ± SD. Statistical significance was tested using one-way ANOVA with multiple comparison tests (F, H). *p* values have been presented in the figure.

**Figure S3.**
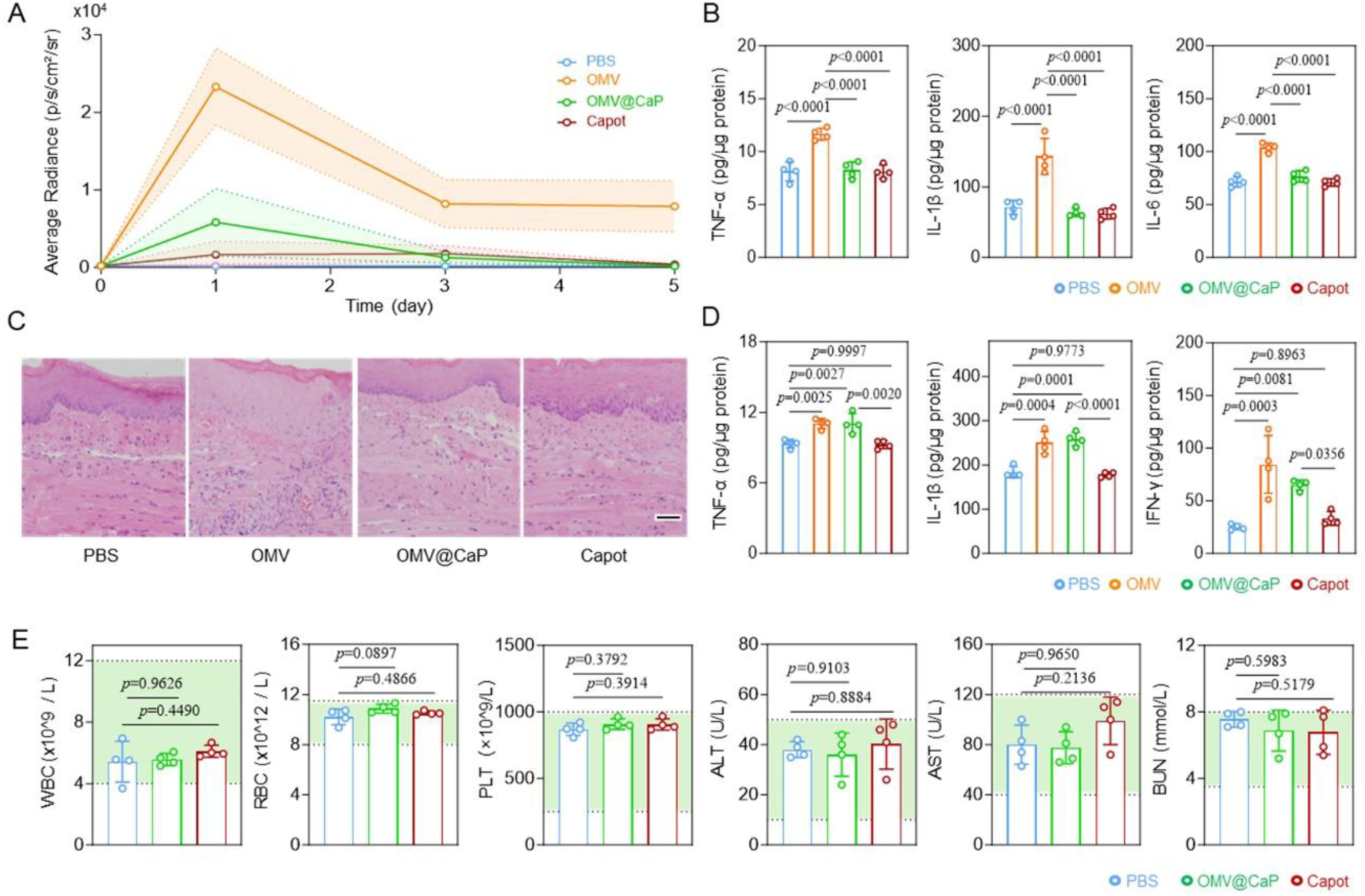
*In vivo* safety evaluations of Capot in mice. A) Corresponding quantitative bioluminescence radiance values for Figure 1K at the indicated time points (n = 4). B) Concentrations of TNF-α, IL-1β, and IL-6 in sublingual tissue at 24 h after sublingual administration with PBS, OMV, OMV@CaP, or Capot (n = 4). C) Representative H&E staining images of ventral tongue tissue at 24 h after sublingual administration with PBS, OMV, OMV@CaP, or Capot. Scale bar = 50 μm. D) Concentrations of TNF-α, IL-1β, and IFN-γ in gastrointestinal tissues at 24 h after sublingual administration with PBS, OMV, OMV@CaP, or Capot (n = 4), which exhibited the original data of Figure 1N. E) Complete blood count (CBC) and blood biochemical (BIOC) test results at 24 h after sublingual administration of PBS, OMV@CaP, or Capot (n = 4). The light green shaded regions denote the normal reference intervals for healthy animals. Data in (A, B, D, E) represent means ± SD. Statistical significance was tested using one-way ANOVA with multiple comparison tests (B, D, E). *p* values have been presented in the figure.

**Figure S4.**
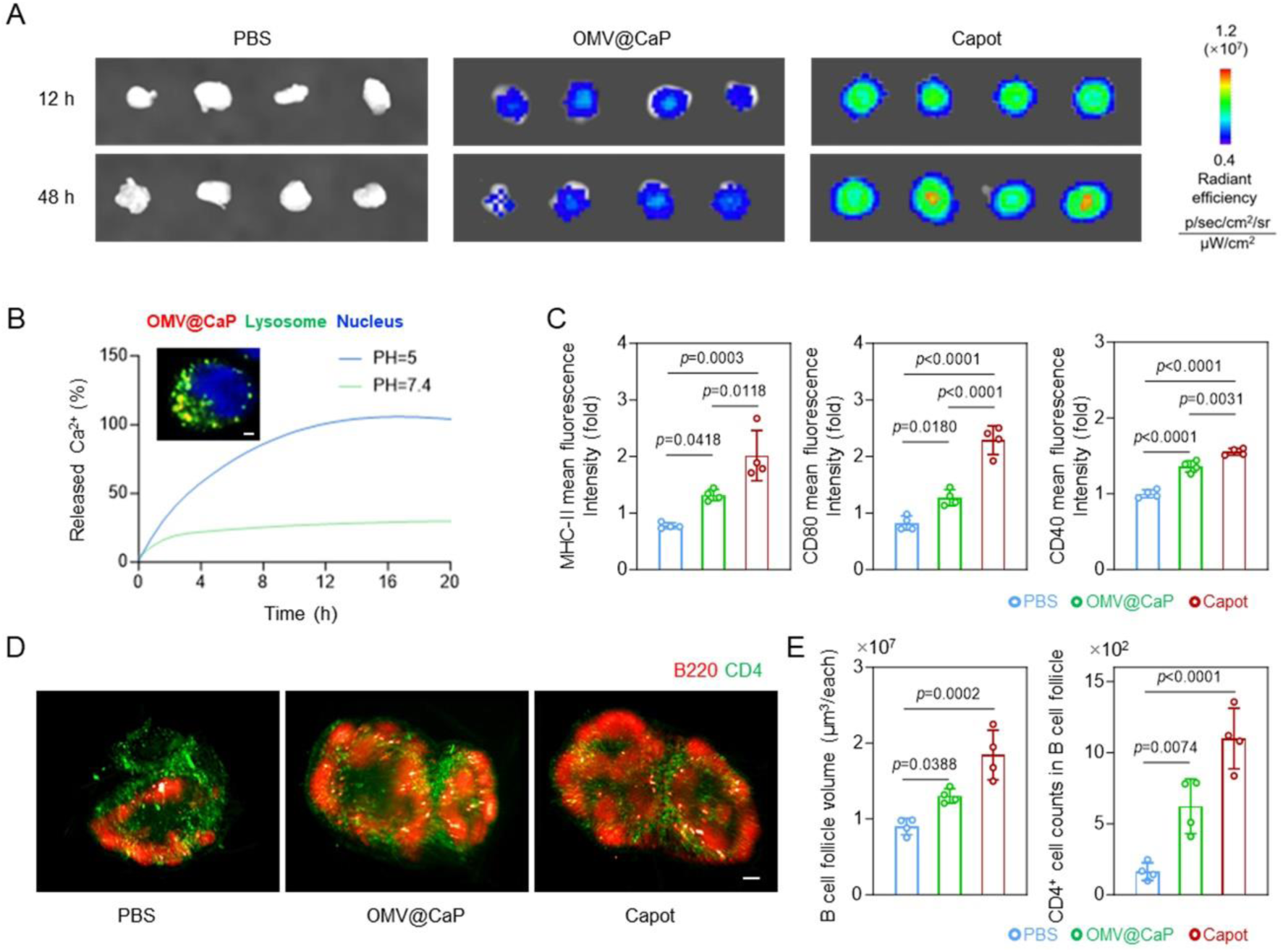
Accumulation and immune activation of Capot in sLN of mice. A) Representative fluorescence images of sLN at indicated time points after sublingual administration of PBS, Cy7-labeled OMV@CaP or Capot. B) Cumulative Ca^2+^ release profiles of OMV@CaPs in normal saline buffer at the indicated pH values, with confocal laser scanning microscopy (CLSM) image showing the colocalization of OMV@CaP and lysosome inside DCs (red, OMV@CaP; green, lysosome; blue, nucleus). Scale bar = 2 μm. C) Flow cytometry analysis showing MHC-II, CD80, and CD40 expression levels on CD11c^+^ DCs in sLNs (n = 4), which exhibited the original data of Figure 2C. D) Light-sheet microscopy images of sLNs (red, B220; green, CD4), which exhibited the original data of Figure 2E. Scale bar = 200 μm. E) Quantification of B cell follicle volume and CD4 T cell infiltration inside B cell follicle of sLN that in (D) (n = 4). Data in (C, E) are presented as means ± SD. Statistical significance was tested using one-way ANOVA with multiple comparison tests (C, E). *p* values have been presented in the figure.

**Figure S5.**
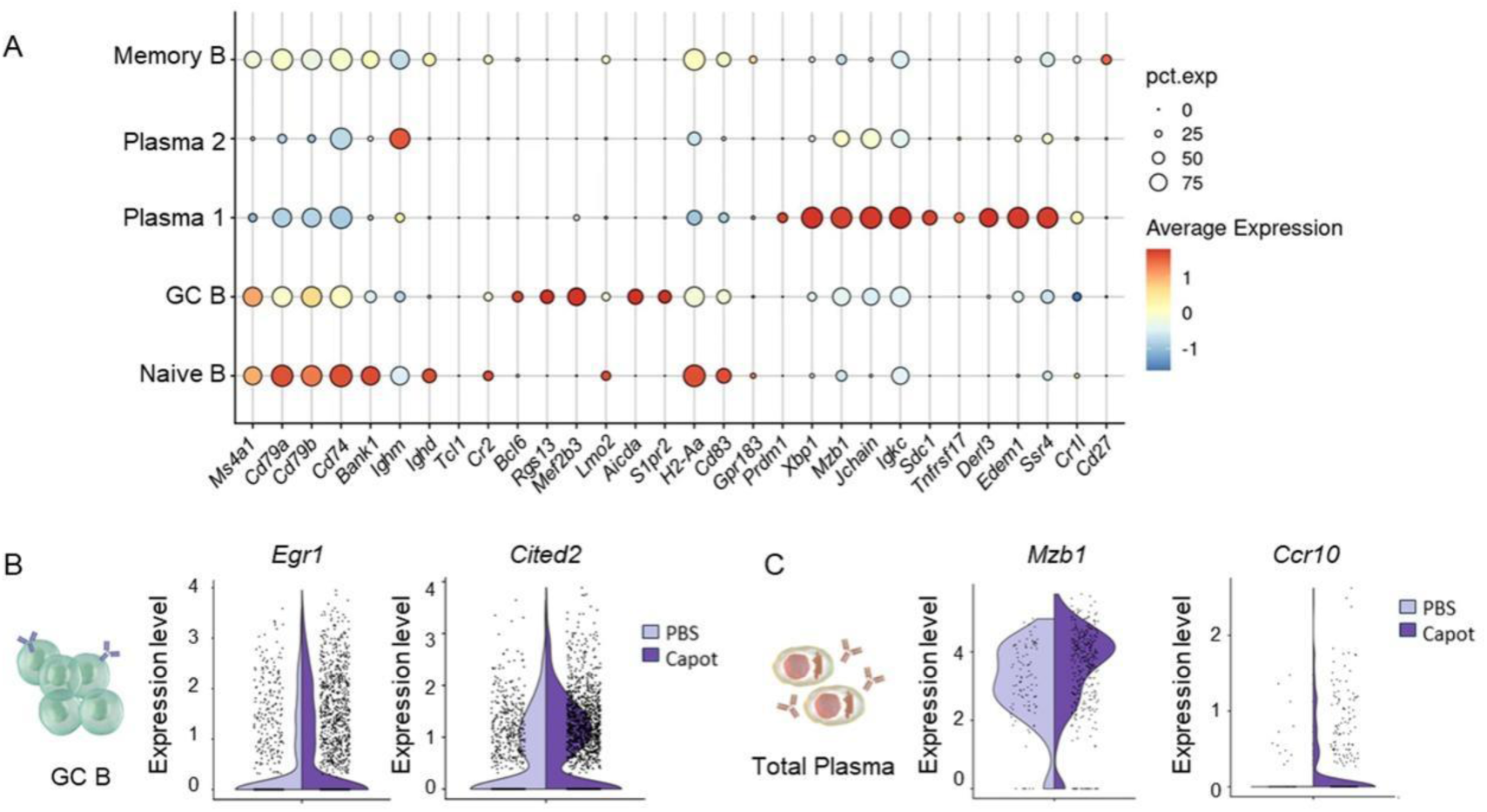
Single-cell sequencing analysis of B cells in sLNs of mice. A) Bubble chart of canonical cell-type markers of five B cell clusters. B) Violin plots comparing the expression of *Egr1* (*p* < 0.0001) and *Cited2* (*p* < 0.0001) in GC B cells between PBS and Capot groups. C) Violin plots comparing the expression of *Mzb1* (*p* = 0.0033) and *Ccr10* (*p* = 0.0008) in Plasma cells between PBS and Capot groups. The width of the violin in (B, C) represents the cell density at different expression values.

**Figure S6.**
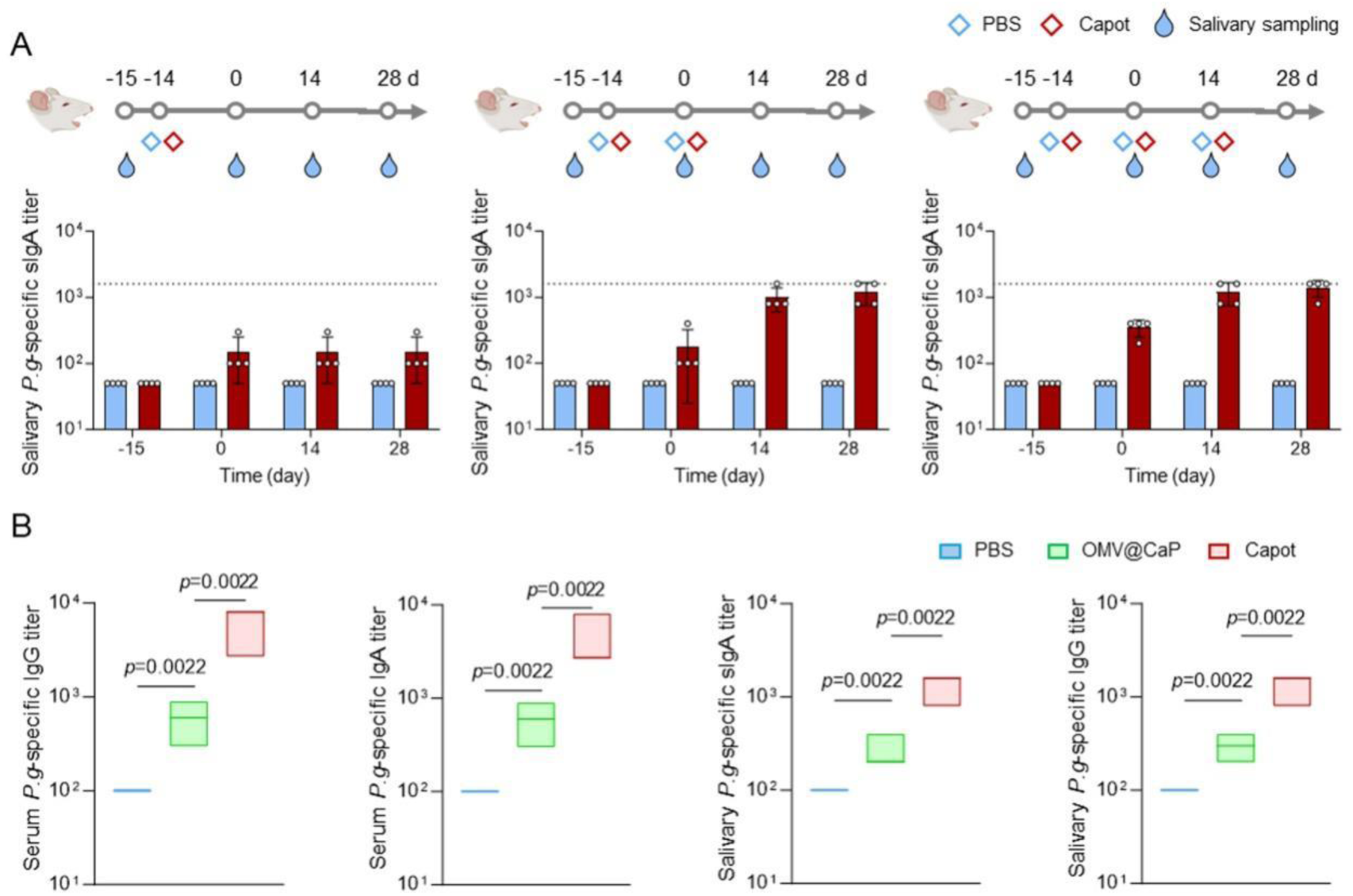
Optimization of the Capot dosing regimen and the corresponding antibody titers in mice. A) Schematic of the immunization and sampling schedule in mice (top) and salivary *P.g*-specific sIgA antibody level under different Capot dosing regimens (bottom, n = 4). B) Serum and salivary *P.g*-specific antibody level at day 14 after twice sublingual administration of PBS, OMV@CaP, or Capot (n = 6). Data in (A) are presented as means ± SD. Box-and-whisker plots in (B) show the median (central line), with the top and bottom of the box representing the maximum and minimum values. Statistical significance was tested using Kruskal-Wallis test with multiple comparison tests in (B). *p* values have been presented in the figure.

**Figure S7.**
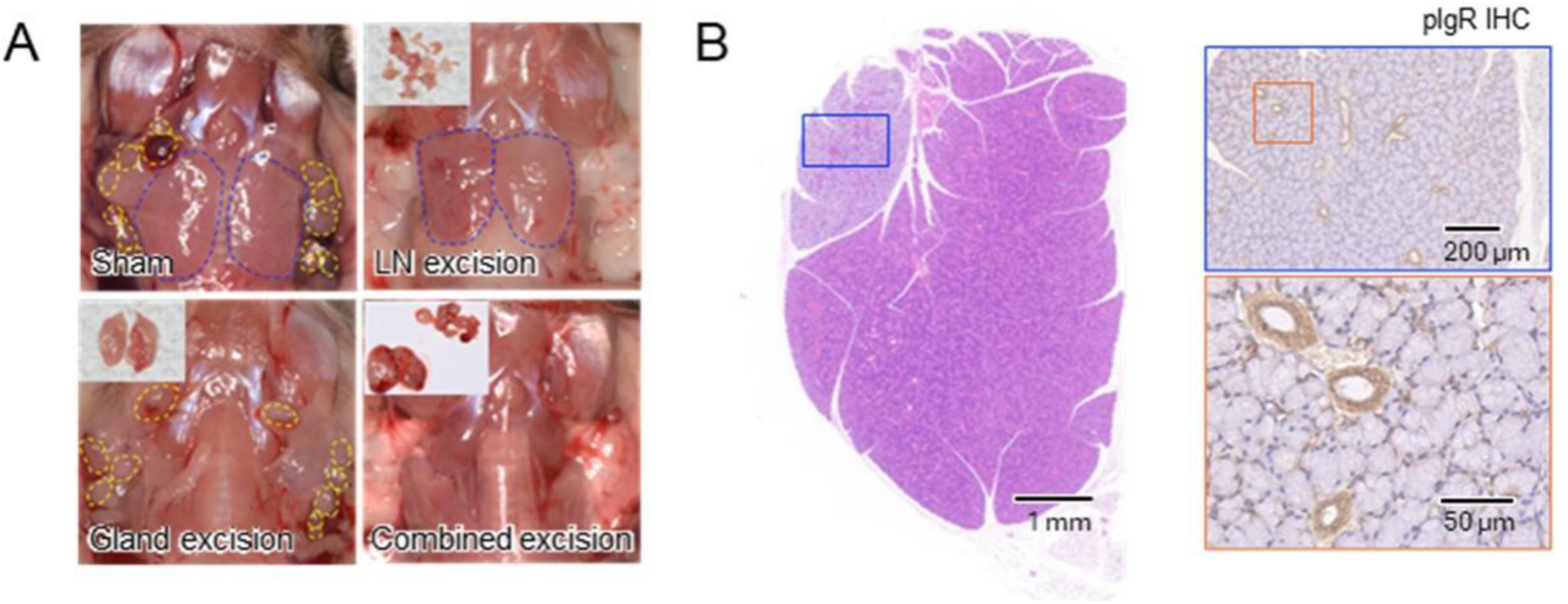
The contribution of sLN and salivary gland in the salivary sIgA secretion. A) Representative photographs showing mice subjected to sham surgery, sLN excision, salivary gland excision, or combined excision. Yellow dashed lines indicate the sLN, and blue dashed lines indicate the gland. B) Representative H&E staining images of salivary gland and corresponding immunohistochemical staining images of pIgR.

**Figure S8.**
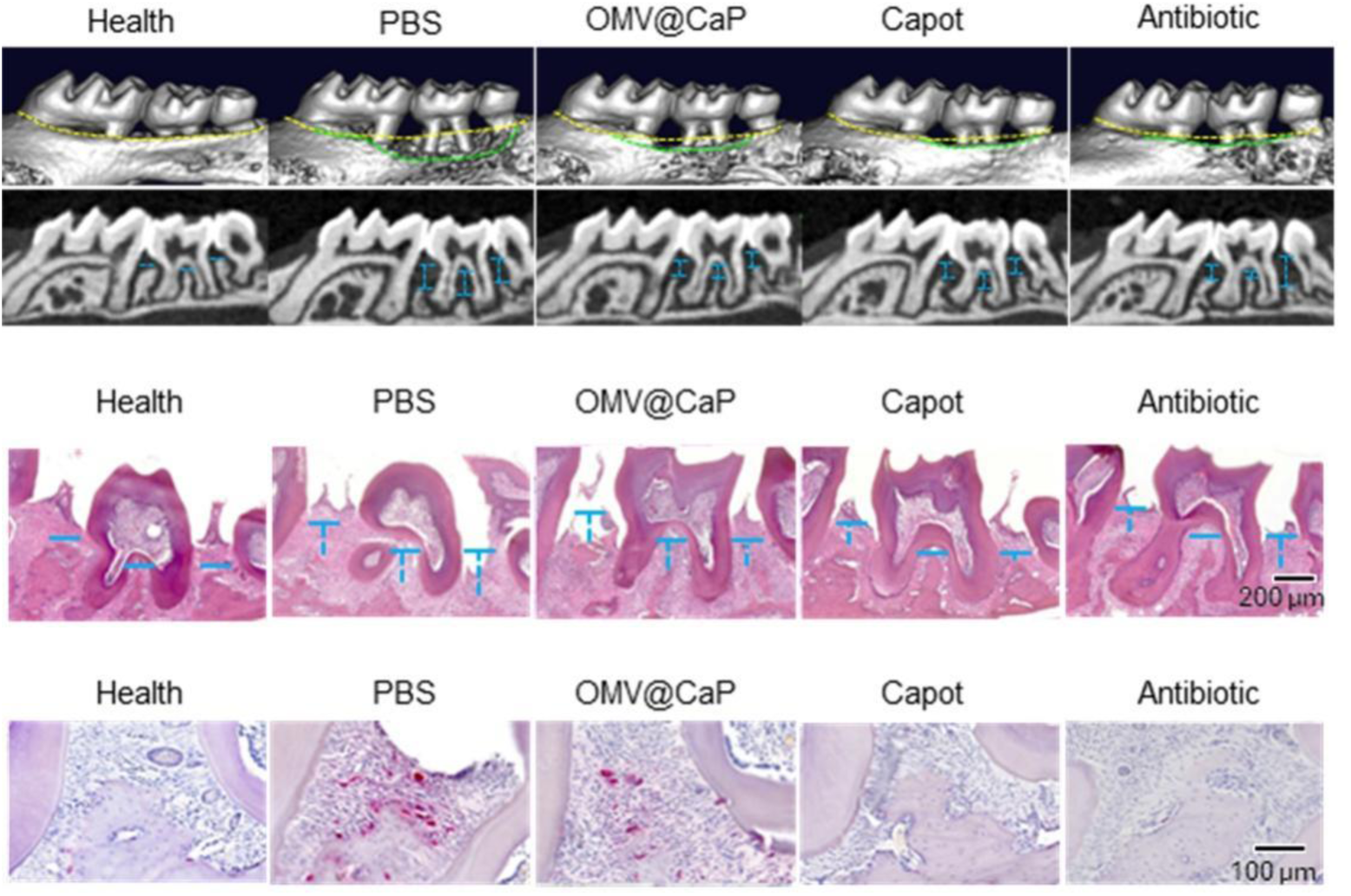
Representative micro-CT (top), H&E staining (middle) and Tartrate-Resistant Acid Phosphatase (TRAP) staining (bottom) images of the experimental tooth in the primary periodontitis model mice that related to Figure 3E. Yellow dashed lines indicate the original height of the alveolar bone in healthy mice. Green dashed lines indicate the height of the alveolar bone in the indicated modeling groups. Blue dashed lines indicate the resorption height of the alveolar bone.

**Figure S9.**
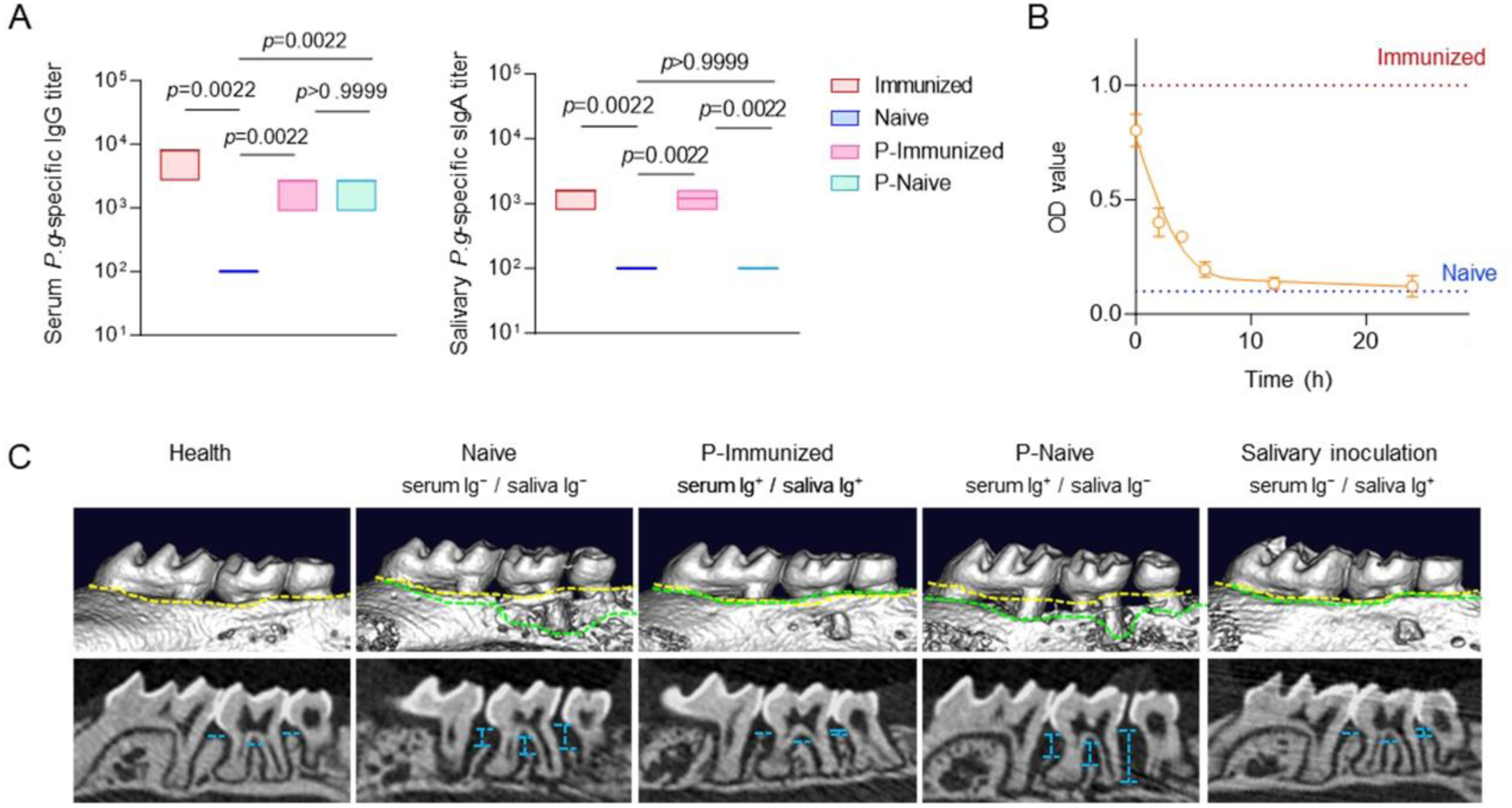
Verification of parabiosis or salivary inoculation models and corresponding evaluation of periodontitis severity. A) Serum and salivary *P.g*-specific antibody level in Capot-immunized mice, naive mice, P-immunized mice of parabiosis model, and P-naive mice of parabiosis model (*n* = 6). B) Attenuation curve of salivary *P.g*-specific antibody in the salivary inoculation model (*n* = 6). C) Representative micro-CT images of the experimental tooth that related to Figure 3F. Yellow dashed lines indicate the original height of the alveolar bone in healthy mice. Green dashed lines indicate the height of the alveolar bone in the indicated modeling groups. Blue dashed lines indicate the resorption height of the alveolar bone. Box-and-whisker plots in (A) show the median (central line), with the top and bottom of the box representing the maximum and minimum values. Data in (B) are presented as means ± SD. Statistical significance was tested using Kruskal-Wallis test with multiple comparison tests in (A). *p* values have been presented in the figure.

**Figure S10.**
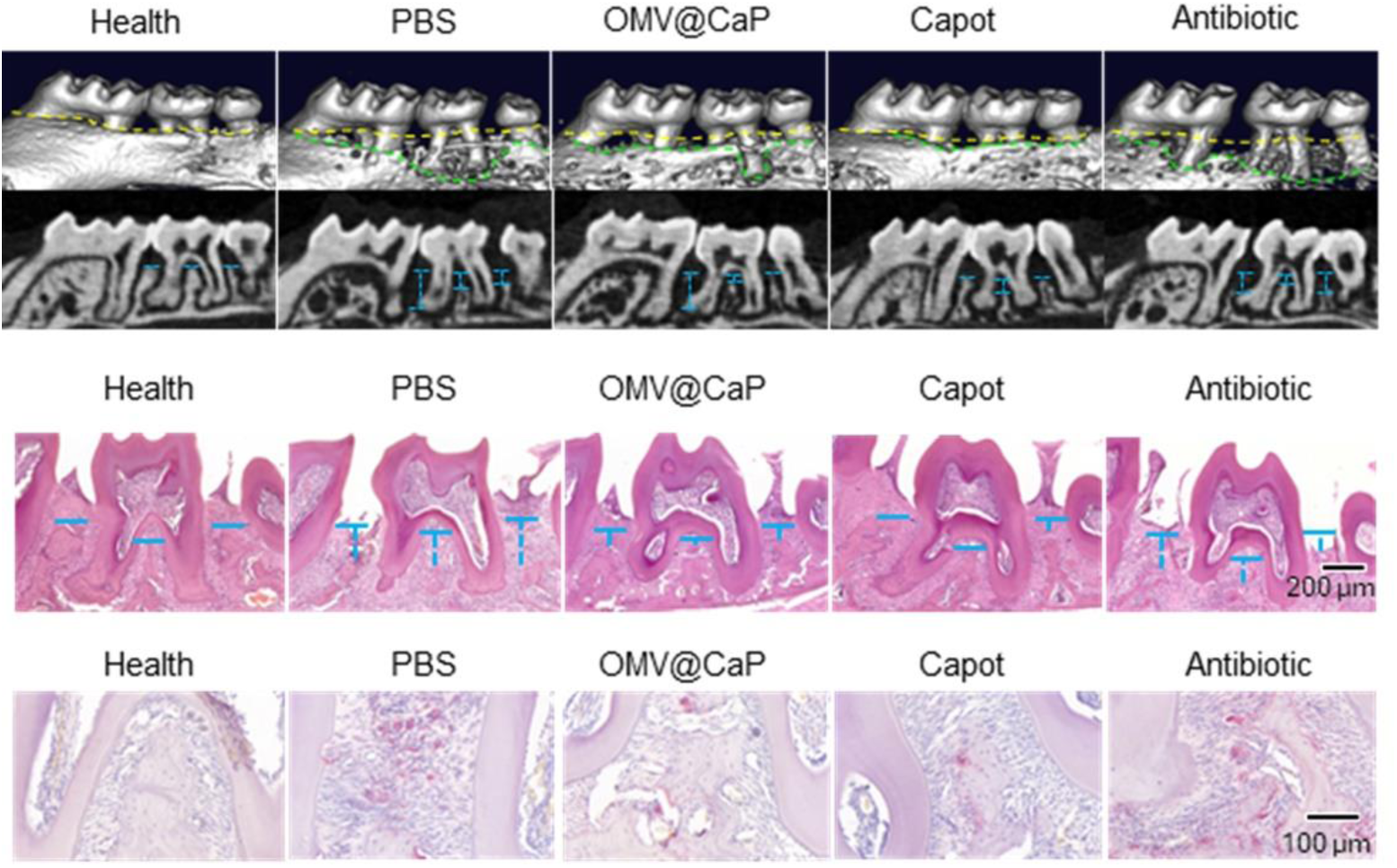
Representative micro-CT (top), H&E staining (middle) and TRAP staining (bottom) images of the experimental tooth in the secondary reinfection periodontitis model mice that related to Figure 3G. Yellow dashed lines indicate the original height of the alveolar bone in healthy mice. Green dashed lines indicate the height of the alveolar bone in the indicated modeling groups. Blue dashed lines indicate the resorption height of the alveolar bone.

**Figure S11.**
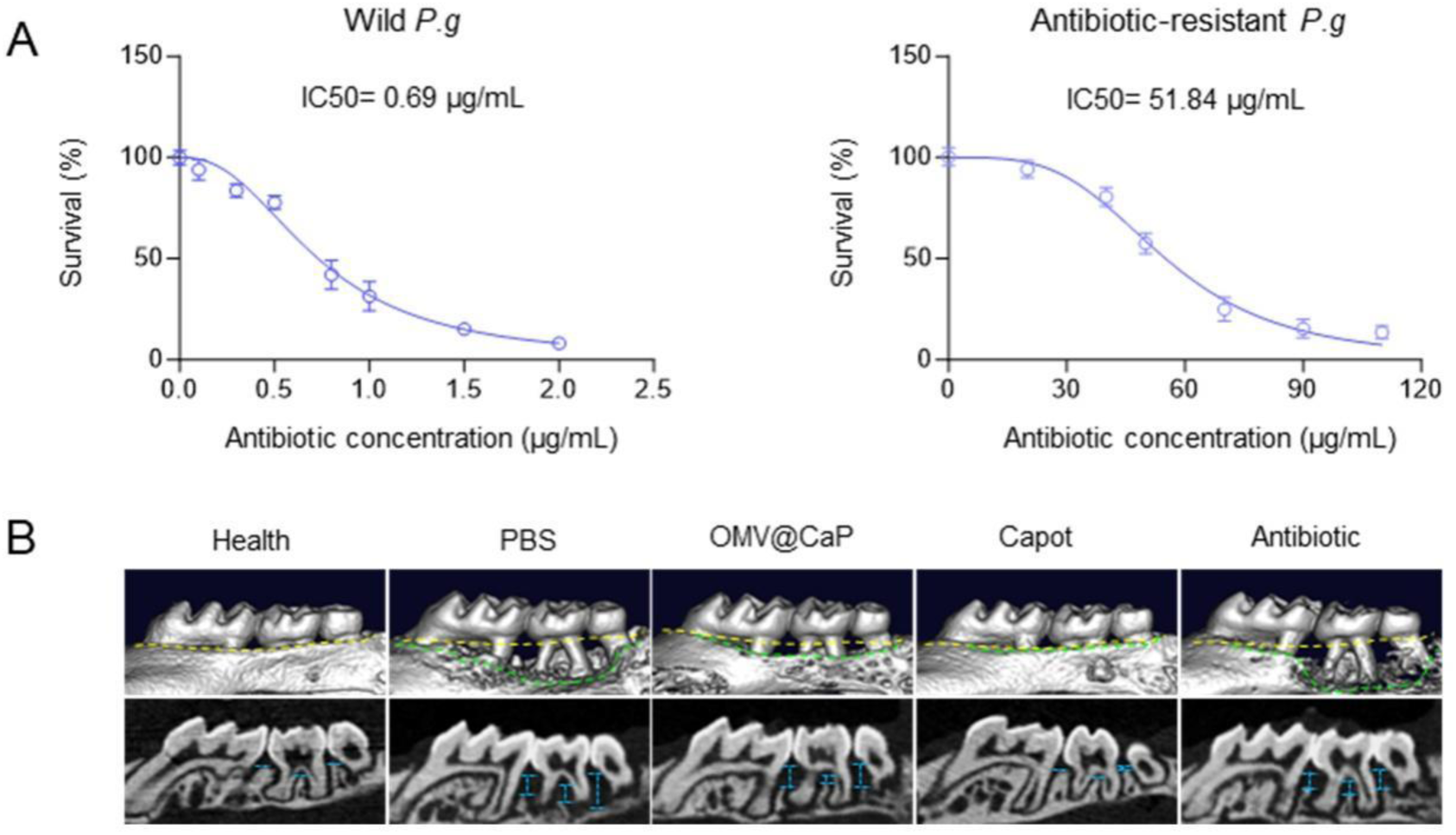
Construction of the antibiotic-resistant *P.g* strain and the periodontitis severity of antibiotic-resistant *P.g*-induced primary periodontitis model. A) Median inhibition concentration curves of wild *P.g* strain and antibiotic-resistant *P.g* strain (n = 6). B) Representative micro-CT images of the experimental tooth that related to Figure 3H. Yellow dashed lines indicate the original height of the alveolar bone in healthy mice. Green dashed lines indicate the height of the alveolar bone in the indicated modeling groups. Blue dashed lines indicate the resorption height of the alveolar bone. Data in (A) are presented as means ± SD.

**Figure S12.**
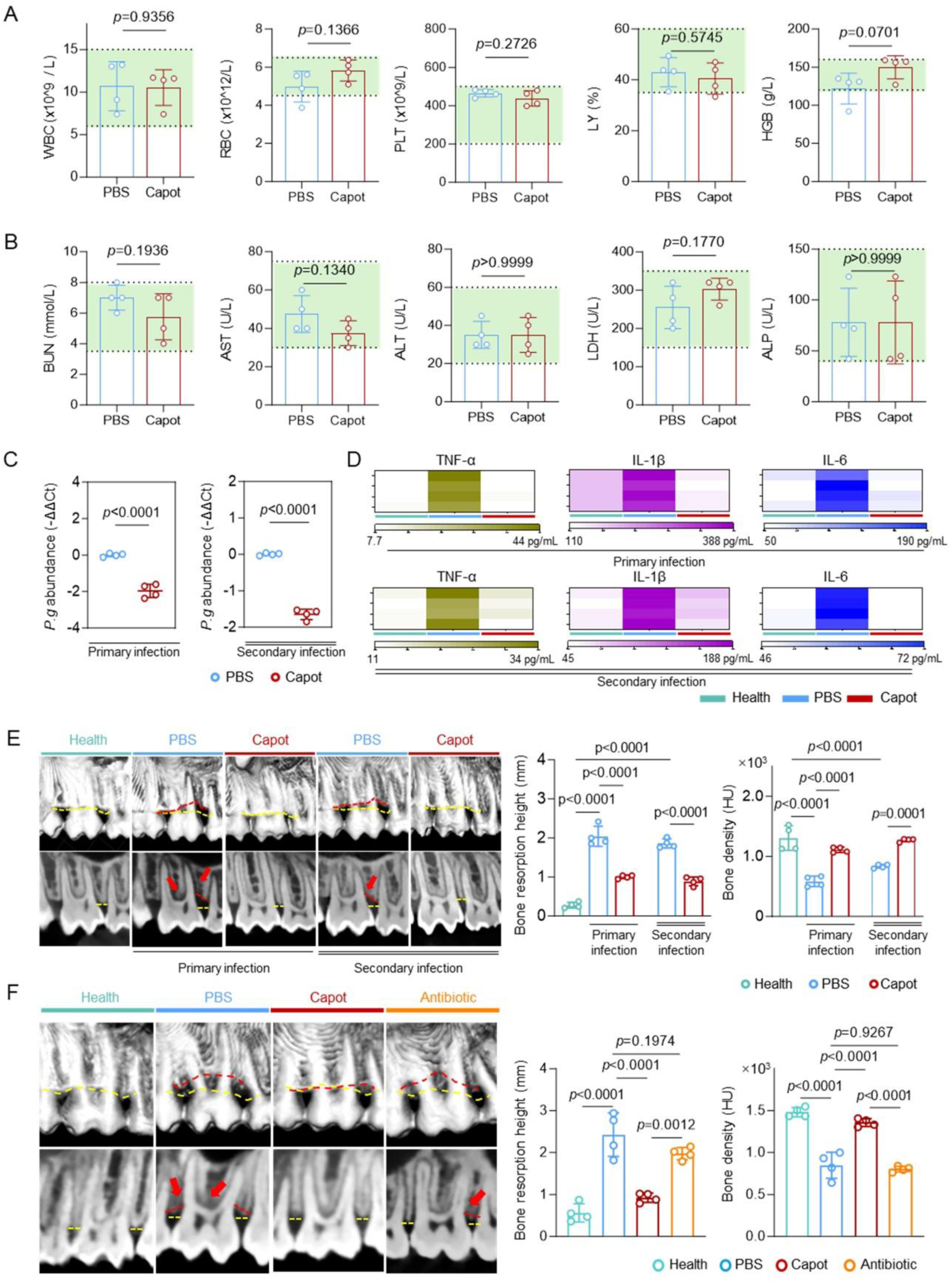
Performance of Capot in cynomolgus monkeys. A) CBC data at 24 h after sublingual administration of PBS or Capot (n = 4). The light green shaded regions denote the normal reference intervals for healthy animals. B) BIOC data at 24 h after sublingual administration of PBS or Capot (n = 4). The light green shaded regions denote the normal reference intervals for healthy animals. C) Quantitative PCR showing salivary *P.g* burden on day 28 (primary infection model; left) and on day 148 (secondary infection model; right) (n = 4). D) Concentrations of salivary TNF-α, IL-1β, and IL-6 on day 28 (primary infection) and day 148 (secondary infection). E) Representative CBCT images (left), and corresponding quantitative analysis including bone resorption height and bone density (right) on day 28 (primary infection) and day 148 (secondary infection) (n = 4). Red arrows indicate areas of alveolar bone destruction. Yellow dashed lines indicate the original height of the alveolar bone in healthy animals. Red dashed lines indicate the height of the alveolar bone in the indicated modeling groups. F) Representative CBCT images (left), and corresponding quantitative analysis including bone resorption height and bone density (right) on day 28 in antibiotic-resistant *P.g* induced periodontitis model (n = 4). Red arrows indicate areas of alveolar bone destruction. Yellow dashed lines indicate the original height of the alveolar bone in healthy animals. Red dashed lines indicate the height of the alveolar bone in the indicated modeling groups. Data in (A-C, E, F) are presented as means ± SD. Statistical significance was tested using unpaired two-tailed t-tests (A-C) and one-way ANOVA with multiple comparison tests (E, F). *p* values have been presented in the figure.

## Notes

### Competing Interest Statement

The authors have declared no competing interest.

